# Structural Covariance Networks in Post-Traumatic Stress Disorder: A Multisite ENIGMA-PGC Study

**DOI:** 10.1101/2021.03.13.432212

**Authors:** Gopalkumar Rakesh, Mark Logue, Emily Clarke-Rubright, Brian M. O’Leary, Courtney C. Haswell, Hong Xie, Paul M. Thompson, Emily L. Dennis, Neda Jahanshad, Saskia B.J. Koch, Jessie L. Frijling, Laura Nawijn, Miranda Olff, Mirjam van Zuiden, Faisal M. Rashid, Xi Zhu, Michael D. De Bellis, Judith K. Daniels, Anika Sierk, Antje Manthey, Jennifer S. Stevens, Tanja Jovanovic, Murray B. Stein, Martha Shenton, Steven J.A. van de Werff, Nic J.A. van de Wee, Robert R.J.M. Vermeiren, Christian Schmahl, Julia Herzog, Milissa L. Kaufman, Lauren O’Connor, Lauren A.M. Lebois, Justin T. Baker, Staci A. Gruber, Jonathan D. Wolff, Erika J. Wolf, Sherry R. Wintemitz, Atilla Gönenc, Kerry J. Ressler, David Bernd Hofmann, Richard A. Bryant, Mayuresh Korgaonkar, Elpiniki Andrew, Li Wang, Ye Zhu, Gen Li, Dan J. Stein, Jonathan Ipser, Sheri Koopowitz, Sven Mueller, Anna Hudson, Luan Phan, Bobak Hosseini, K. Mike Angstadt, Anthony P. King, Marijo Tamburrino, Brynn C. Skilliter, Elbert Geuze, Sanne J.H. van Rooij, Tim Varkevisser, Katie A. McLaughlin, Margaret A. Sheridan, Matthew Peverill, Kelly Sambrook, Dick J. Veltman, Kathleen Thomaes, Steven M. Nelson, Geoffrey May, Lee Baugh, Gina Forster, Raluca Simons, Jeffrey Simons, Vincent Magnotta, Kelene A Fercho, Adi Maron-Katz, Stefan du Plessis, Seth Disner, Nicholas Davenport, Sophia I. Thomopoulos, Benjamin Suarez-Jimenez, Tor D. Wager, Yuval Neria, Negar Fani, Henrik Walter, Inga Koerte, Jessica Bomyea, Kyle Choi, Alan N. Simmons, Elizabeth Olson, Isabelle Rosso, Thomas Straube, Theo G.M. van Erp, Tian Chen, Andrew S. Cotton, John Wall, Richard J. Davidson, Terri deRoon-Cassini, Jacklynn Fitzgerald, Christine Larson, Evan Gordon, Dan Grupe, Scott R. Sponheim, Amit Etkin, Soraya Seedat, Ilan Harpaz-Rotem, Kristen Wrocklage, Chadi G. Abdallah, John H. Krystal, Ifat Levy, Hassaan Gomaa, Mary Agnes B. McMahon, Israel Liberzon, Xin Wang, Delin Sun, Rajendra A. Morey

**Affiliations:** Duke-UNC Brain Imaging and Analysis Center (BIAC); Mid-Atlantic Mental Illness Research, Education and Clinical Center, Durham VA Medical Center, Durham, NC, USA; National Center for PTSD, VA Boston Healthcare System, Boston, MA, USA; Biomedical Genetics, Boston University, Boston, MA, USA; Department of Electrical Engineering and Computer Science, University of Toledo, Toledo, OH, USA; Department of Neurosciences, University of Toledo, Toledo, OH, USA; Imaging Genetics Center, Mark and Mary Stevens Neuroimaging and Informatics Institute, Keck School of Medicine of University of Southern California, Marina del Rey, CA, USA; TBI and Concussion Center, Department of Neurology, University of Utah School of Medicine, Salt Lake City, UT, USA; Academic Medical Center, Amsterdam, Netherlands; Donders Center for Cognitive Neuroimaging, Radboud University Nijmegen, Nijmegen, Netherlands; VU University Medical Center, Amsterdam, the Netherlands; ARQ National Psychotrauma Centre, Diemen, The Netherlands; Amsterdam Neuroscience & Amsterdam Public Health Research Institutes, Amsterdam, the Netherlands; Columbia University, New York, NY, USA; University of Groningen, Groningen, Netherlands; University Medical Center Charite, Berlin, Germany; Department of Psychiatry and Behavioral Sciences, Emory University, Atlanta, GA, USA; Department of Psychiatry, University of California, San Diego, La Jolla, CA, USA; Harvard University, Boston, MA, USA; Leiden University Medical Center, Leiden, Netherlands; Department of Psychosomatic Medicine and Psychotherapy, Central Institute of Mental Health Mannheim, Medical Faculty Mannheim, Heidelberg University, Mannheim, Germany; Department of Psychiatry, University of Western Ontario, London, ON, Canada; McLean Hospital, Harvard University, Boston, MA, USA; CUNY Graduate Center, New York, NY, USA; Department of Psychiatry, Harvard Medical School, Boston, MA, USA; Department of Counseling, Developmental, and Educational Psychology, Boston, MA, USA; Department of Psychiatry, Boston University School of Medicine, Boston, MA, USA; University Hospital Münster, Germany; University of New South Wales, Sydney, Australia; Brain Dynamics Centre, Westmead Institute for Medical Research, University of Sydney, Sydney, Australia; University of Sydney, Sydney, Australia; Institute of Psychology, Chinese Academy of Sciences, Beijing, China; University of Cape Town, Cape Town, South Africa; Department of Experimental Clinical and Health Psychology, Ghent University, Henri Dunantlaan 2, 9000 Ghent, Belgium; Department of Personality, Psychological Assessment and Treatment, University of Deusto, Bilbao, Spain; The Ohio State University Wexner Medical Center, Columbus, OH, USA; Department of Psychiatry, University of Illinois at Chicago, Chicago, IL, USA; Mental Health Service Line, Jesse Brown VA Medical Center, Chicago, IL, USA; Department of Psychiatry and Behavioral Science, Texas A&M Health Science Center, College of Medicine, College Station, TX, USA; University of Michigan, Ann Arbor, MI, USA; Department of Psychiatry, University of Toledo, Toledo, OH, USA; University Medical Center Utrecht, Utrecht, The Netherlands; University of North Carolina, Chapel Hill, NC, USA; University of Washington, Seattle, WA, USA; VU University Medical Center, Amsterdam, Netherlands; Sinai Centrum and VU University Medical Center, Amsterdam, Netherlands; VISN 17 Center of Excellence for Research on Returning War Veterans, Waco, TX, USA; Center for Vital Longevity, University of Texas at Dallas, Dallas, TX, USA; Division of Basic Biomedical Sciences, Sanford School of Medicine, University of South Dakota, Vermillion, SD, USA; Brain Health Research Centre, Department of Anatomy, University of Otago, Dunedin, New Zealand; Center for Brain and Behavior Research, University of South Dakota, Vermillion, SD, USA. 20; Department of Psychology, University of South Dakota, Vermillion, SD, USA; Departments of Radiology, Psychiatry, and Biomedical Engineering, University of Iowa, Iowa City, IA, USA; Civil Aerospace Medical Institute, US Federal Aviation Administration, Oklahoma City, OK, USA; Stanford University, Palo Alto, CA, USA; Stellenbosch University, Stellenbosch, South Africa; Minneapolis VA Health Care System, Minneapolis, MN, USA; Department of Psychiatry, University of Minnesota, Minneapolis, MN, USA; Dartmouth College, Hanover, NH, USA; Psychiatry Neuroimaging Laboratory, Brigham & Women’s Hospital, Boston, MA; Institute of Medical Psychology and Systems Neuroscience, University of Münster; Clinical Translational Neuroscience Laboratory, Department of Psychiatry and Human Behavior, University of California Irvine, Irvine, CA, USA, 92617, USA; Department of Mathematics and Statistics, University of Toledo, Toledo, OH, USA; University of Wisconsin-Madison; Department of Surgery, Division of Trauma & Acute Care Surgery; Comprehensive Injury Center, Medical College of Wisconsin; Department of Psychology, Marquette University; Department of Psychology, University of Wisconsin-Milwaukee; Department of Radiology, Washington University School of Medicine, St. Louis, MO, USA; Clinical Neuroscience Division, VA National Center for PTSD, VA Connecticut Healthcare System, New Haven, CT, USA; Yale University School of Medicine, New Haven, CT, USA; Michael E. DeBakey, VA Medical Center, Houston, TX, USA; Menninger Department of Psychiatry, Baylor College of Medicine, Houston, TX, USA; Department of Psychiatry and Behavioral Health, Penn State College of Medicine

## Abstract

**Introduction:** Cortical thickness (CT) and surface area (SA) are established biomarkers of brain pathology in posttraumatic stress disorder (PTSD). Structural covariance networks (SCN) constructed from CT and SA may represent developmental associations, or unique interactions between brain regions, possibly influenced by a common causal antecedent. The ENIGMA-PGC PTSD Working Group aggregated PTSD and control subjects’ data from 29 cohorts in five countries (n=3439).

**Methods:** Using Destrieux Atlas, we built SCNs and compared centrality measures between PTSD subjects and controls. Centrality is a graph theory measure derived using SCN.

**Results:** Notable nodes with higher CT-based centrality in PTSD compared to controls were left fusiform gyrus, left superior temporal gyrus, and right inferior temporal gyrus. We found sex-based centrality differences in bilateral frontal lobe regions, left anterior cingulate, left superior occipital cortex and right ventromedial prefrontal cortex (vmPFC). Comorbid PTSD and MDD showed higher CT-based centrality in the right anterior cingulate gyrus, right parahippocampal gyrus and lower SA-based centrality in left insular gyrus.

**Conclusion:** Unlike previous studies with smaller sample sizes (≤318), our study found differences in centrality measures using a sample size of 3439 subjects. This is the first cross-sectional study to examine SCN interactions with age, sex, and comorbid MDD. Although limited to group level inferences, centrality measures offer insights into a node’s relationship to the entire functional connectome unlike approaches like seed-based connectivity or independent component analysis. Nodes having higher centrality have greater structural or functional connections, lending them invaluable for translational treatments like neuromodulation.

## 1. INTRODUCTION

Post-traumatic stress disorder (PTSD) has a lifetime prevalence of 9.4% among adults in the US (Kessler et al., 2005) and 4% globally (Liu et al., 2017). Cross-sectional and longitudinal studies show structural changes to specific brain regions and structural and functional connectivity differences between regions in PTSD (Akiki, Averill, & Abdallah, 2017; Hughes & Shin, 2011; Mueller et al., 2015; Philip, Carpenter, & Sweet, 2014; Tursich et al., 2015). Cortical thickness (CT) and surface area (SA) are reliable biomarkers of pathology across psychiatric illnesses including PTSD. Interregional relationships in cortical thickness (Yun et al., 2020) are referred to as *structural covariance network*s (SCN). Features of a SCN, such as centrality, may be used to characterize regional and network pathology associated with neuropsychiatric disorders. Centrality is a concept from graph theory, which measures the importance of a particular region within a network. In graph theory, a network is made up of connections (edges) between brain regions (nodes)(Rubinov & Sporns, 2010b). Cortical volume, CT, and SA have been used to generate centrality measures using all possible pairwise correlations between cortical regions as edges in a group of subjects (He & Evans, 2010; Rubinov & Sporns, 2010a; Sun, Haswell, Morey, & De Bellis, 2018; Sun, Peverill, Swanson, McLaughlin, & Morey, 2018). SCNs that characterize interregional correlations of CT or SA in group of subjects may represent underlying functional associations (Gong, He, Chen, & Evans, 2012; He, Chen, & Evans, 2007). Positive interregional correlations based on CT are consistent with diffusion imaging derived structural connections (Gong et al., 2012) and genome co-expression (Romero-Garcia et al., 2018).

Studies have investigated PTSD-associated differences in cortical measures (Bromis, Calem, Reinders, Williams, & Kempton, 2018; Wang et al., 2020), task-based functional connectivity (Hughes & Shin, 2011), and resting-state functional connectivity (Koch et al., 2016). Thus far, meta-analyses of structural neuroimaging in PTSD have applied voxel-based morphometry (VBM) and volume estimates of cortical regions to reveal gray matter volume differences in anterior cingulate cortex, insula, medial and ventromedial prefrontal cortex, orbitofrontal cortex, left temporal pole, rostral middle frontal gyrus, and superior frontal gyrus (Kuhn & Gallinat, 2013; Meng et al., 2016). However, there has been sparse literature on SCN in PTSD, consisting of two studies in adults with PTSD (Mueller et al., 2015; Sun, Davis, et al., 2018), two studies examining SCN in children or youth with PTSD (Sun, Haswell, et al., 2018; Sun, Peverill, et al., 2018), one study focused on SCN derived from diffusion tensor imaging (DTI) measures (Long et al., 2013), and finally one longitudinal study focused on SCN features predicting symptom onset following acute exposure to trauma (Harnett NG, Available online 7 August 2020). Findings were not consistent across studies, areas highlighted included bilateral anterior cingulate, bilateral superior frontal gyrus, right insula and occipital cortex. The sample sizes of the aforementioned studies were relatively small (~300) in relation to the 148 cortical regions under consideration, which involves 10,878 inter-regional relationships.

High data dimensionality in relation to sample size presents a challenge in controlling Type I error (Konietschke, Schwab, & Pauly, 2020). Thus, drawing robust inferences from a high-dimensional feature space containing 148 cortical regions (Destrieux Atlas) requires a sample size that is larger than the number of features. Furthermore, each of the published studies have focused on specific types of trauma and/or populations such as military (Mueller et al., 2015), motor vehicle crash (Harnett NG, Available online 7 August 2020), childhood maltreatment (Sun, Haswell, et al., 2018; Sun, Peverill, et al., 2018), but there is lack of SCN results that capture the heterogeneity of trauma types commonly found in PTSD. Finally, none of these published reports in adults with PTSD examined cortical surface area derived SCNs. Surface area is of particular interest given its strong genetic basis in relation to the weaker genetic basis of cortical thickness. A large GWAS examining the genetic architecture of the human cortex found that surface area was negatively genetically correlated to cortical thickness (Grasby et al., 2020). Given the divergent genetic architectures of surface area and cortical thickness coupled with the role of genetics to PTSD (Nievergelt et al., 2019) means it is essential to dissect the distinct contributions of CT-derived and SA-derived SCNs.

There have been no studies to investigate sex differences in PTSD using SCN. In the present study, we compared CT- and SA-based network centrality measures in 3439 PTSD and trauma-exposed control subjects from the Enhancing Neuroimaging Genetics through Meta-Analysis (ENIGMA) - Psychiatric Genomics Consortium (PGC) Consortium for PTSD. We hypothesized that areas previously implicated in the pathophysiology of PTSD such as the anterior cingulate, insula, ventromedial prefrontal cortex (vMPFC) and occipital regions would show differences in network centrality. Given sex-based difference in salience network connectivity in PTSD, we hypothesized sex-based disruptions in network centrality measures in the salience network comprising dorsal anterior cingulate (dACC), and insula. Consistent with limited previous literature, we also hypothesized centrality differences in the parahippocampal gyrus and salience network based on presence or absence of comorbid major depressive disorder (MDD).

## 2. SUBJECTS AND METHODS

### 2.1 Subjects

The ENIGMA-PGC PTSD Working Group aggregated (Table 1) PTSD patients and control subjects’ data with varying levels of trauma exposure from 29 cohorts in five countries. We analyzed CT data from 3439 subjects and SA data from 3436 subjects. The vast majority (92% of subjects with PTSD, 85% of control subjects) of participants were adults, and the remaining sample was a combination of children and adolescents (Table 1). The sample comprised 1348 PTSD subjects (39%) and 2082 healthy controls (61%). Average age and standard deviation (SD) of male PTSD subjects was (38±15) years, (37±17.1) years for male control subjects, (34.5± 12.7) years for female PTSD subjects, (32±14.3) years for female control subjects. **Supplemental table 1** shows age by diagnoses and sex at each site. Sixteen out of 29 sites had control subjects with trauma exposure (**Table 1**); the remaining sites had controls who were unexposed to trauma. The percentage of subjects with MDD is also reported in **Table 1**.

### 2.2 Methods

Rating scales are described in section 2.2.1 of supplementary material.

Scanner details and acquisition parameters are in **Supplementary Table 2**. Inclusion and exclusion criteria for each site are in **Supplementary Table 24**. All participating sites obtained approval from local institutional review boards and ethics committees. All participants provided written informed consent.

#### 2.2.2 Imaging

Structural MRI (T1) data obtained from cross-sectional case-control studies were analyzed at Duke University with a standardized neuroimaging and QC pipeline developed by the ENIGMA Consortium (http://enigma.ini.usc.edu/protocols/imaging-protocols/) (Logue et al., 2018). Cortical parcellation was performed with FreeSurfer 5.3 on 148 regions (74 per hemisphere) that were labeled with the Destrieux atlas (Destrieux, Fischl, Dale, & Halgren, 2010).

#### 2.2.3 Network analyses

#### 2.3.6 Centrality measures

We selected network topology measures that would enable direct comparisons of our results with previous findings (Mueller et al., 2015; Teicher, Anderson, Ohashi, & Polcari, 2014). Using undirected connections for correlation-derived graphs, four centrality measures were calculated with the Brain Connectivity Toolbox (BCT) (Rubinov & Sporns, 2010a): degree centrality, betweenness centrality, closeness centrality, and eigenvector centrality (**Supplementary section 2.2.6**).

#### 2.2.7 Statistical Analysis

Analyses were performed with MATLAB scripts that we reported previously (Sun, Haswell, et al., 2018; Sun, Peverill, et al., 2018). Our method is similar to, but far more stringent than, methods for controlling Type 1 error used in earlier reports (Teicher et al., 2014). Further details are available in the **Supplementary material 2.2.7**.

## 3. RESULTS

Key demographic and clinical characteristics of each cohort are reported in **Table 1**. Wiring costs obtained from CT-based positive correlation coefficients were 0.06 and 0.07 for PTSD and control groups respectively, and SA-based wiring costs were 0.1 and 0.007 in PTSD and control groups respectively. These minimum wiring costs were calculated using supra-threshold positive correlation and ensured that nodes in the network were well connected (Teicher et al., 2014). **Table 3** shows wiring costs for each diagnostic group.

### 3.1 CT SCN Results

#### 3.1.1 Graph Centrality Based on Pearson’s Correlation Coefficients

The PTSD group showed higher centrality compared to controls in the left fusiform gyrus, left precentral gyrus, lateral aspect of left superior temporal gyrus, left inferior frontal sulcus, right paracentral gyrus and sulcus, right inferior temporal gyrus. The PTSD group showed lower centrality compared to controls in right posterior transverse collateral sulcus and right anterior occipital sulcus (**Table 4 and figure 2**).

**Figure 1.**
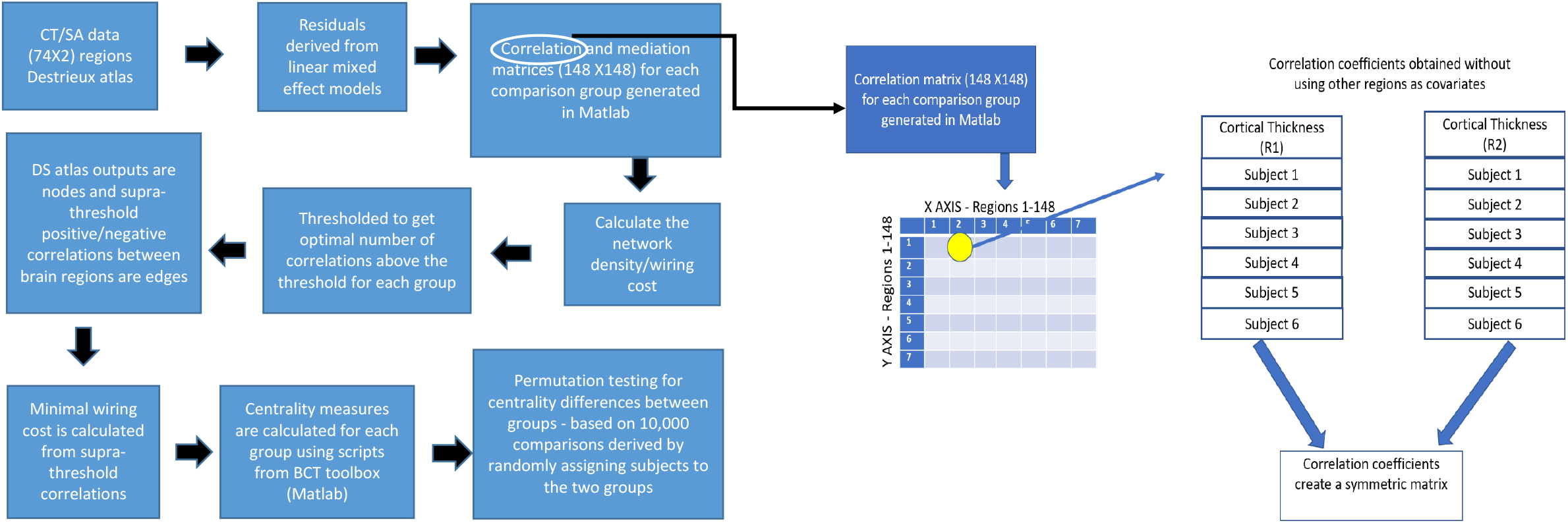
We harmonized CT and SA values with *ComBat* to minimize sources of variance related to study site and scanner while preserving signal associated with the parameters of interest such as diagnostic grouping and other clinical covariates of interest (Fortin et al., 2018). as detailed in supplementary section 2.2.4. Subsequently, age, age^2^, sex, site and mean whole-brain CT or SA estimates were regressed from the CT and SA estimates with a linear model (He et al., 2007) as explained in supplementary section 2.2.5. We performed SCN analyses by representing brain regions as nodes and CT/SA correlation coefficients as edges (He & Evans, 2010; Rubinov & Sporns, 2010a; Sun, Haswell, et al., 2018). Positive and negative coefficients were used to generate separate networks. Wiring cost was used to generate binary graphs for each group and is defined as the number of edges present divided by maximum possible number of edges (Bassett et al., 2008; Teicher et al., 2014). Previous publications considered only positive correlations between regions in centrality analyses as substantiated by evidence that they are largely mediated by direct fiber pathways (Gong et al., 2012). We focused on positive correlations in this report but results of negative correlations are available in the **Supplementary Material**. We assessed the reliability (99% confidence interval) of between-group comparisons using jackknife resampling. Permutation testing generated the probability of significant between-group differences in centrality measures based on the between-group difference of nodal centrality based on a distribution 10,000 random assignments of group. We tested interactions with permutation testing with some modifications of the procedure.

**Figure 2.**
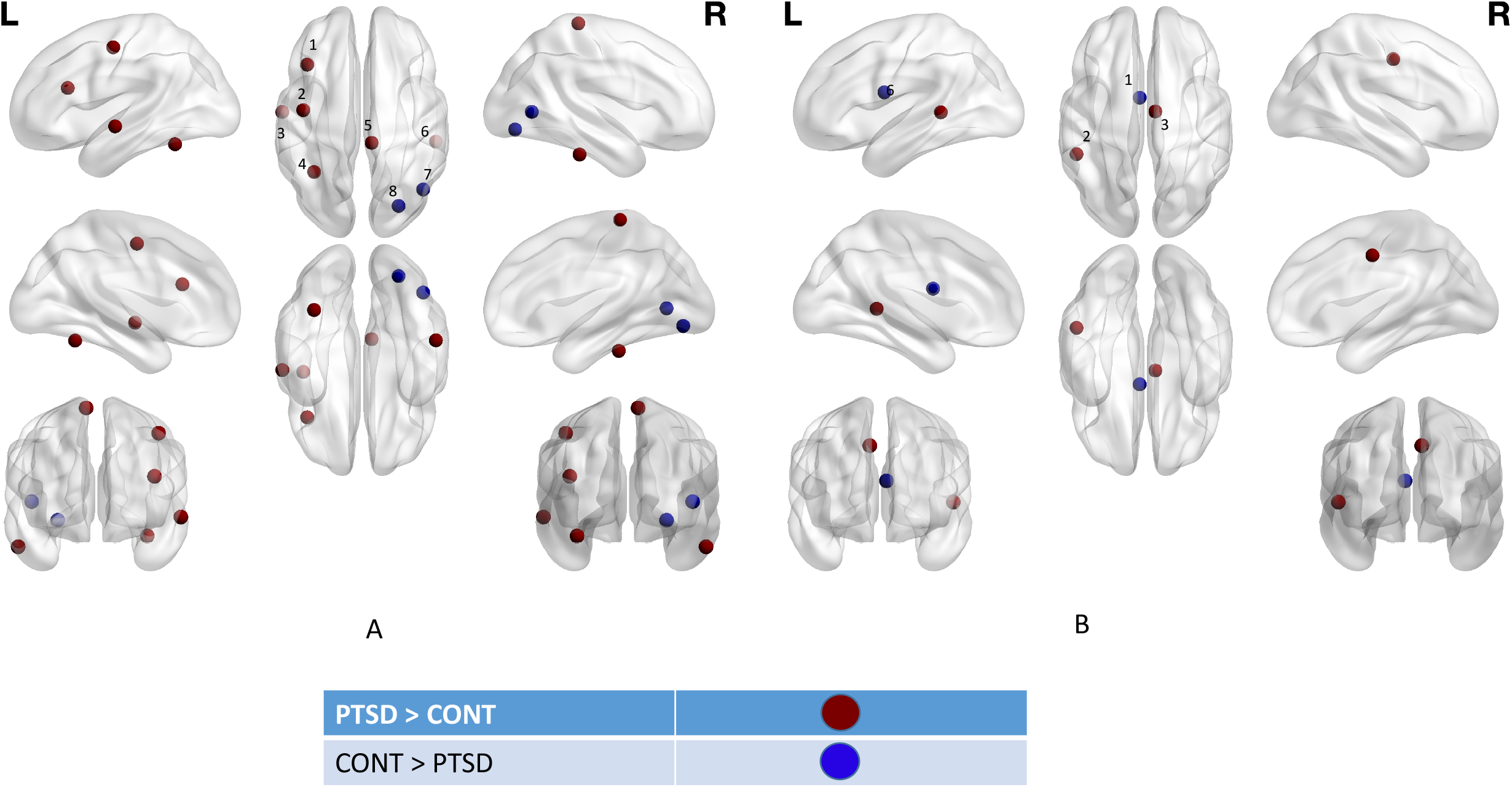
shows centrality differences between PTSD subjects and controls using networks, created only with positive correlations as edges. A highlight these differences with cortical thickness (CT) and B highlights these differences with surface area (SA). For CT, areas corresponding to numbered regions are as follows: - 1-left inferior frontal sulcus, 2-left precentral gyrus, 3-anterior aspect of superior temporal gyrus, 4-left fusiform gyrus, 5-right paracentral gyrus and sulcus, 6– right inferior temporal gyrus, 7-right posterior transverse collateral sulcus, 8– right anterior occipital sulcus. For SA, areas corresponding to numbered regions are as follows: - 1-left pericallosal sulcus, 2-left superior temporal sulcus, 3-right middle posterior part of the cingulate gyrus and sulcus. Regions colored in red showed higher centrality in PTSD subjects compared to controls. Regions colored in blue showed higher centrality in controls compared to PTSD subjects.

**Supplemental Table 4** provides comparisons of centrality between PTSD and control groups based on negative correlation coefficients.

#### 3.1.2 Interactions

Brain regions with a significant interaction of age and diagnosis on centrality that are based on positive correlation coefficients are displayed in Figure 3 and listed in **Supplemental Tables 5** to **12** for age ranges (in years) <10, 10-15, 15-20, 20-30, 30-40, 40-50, 50-60, and >60 respectively. **Supplemental figure 1** shows the number of subjects in each age group for PTSD subjects versus controls.

**Figure 3.**
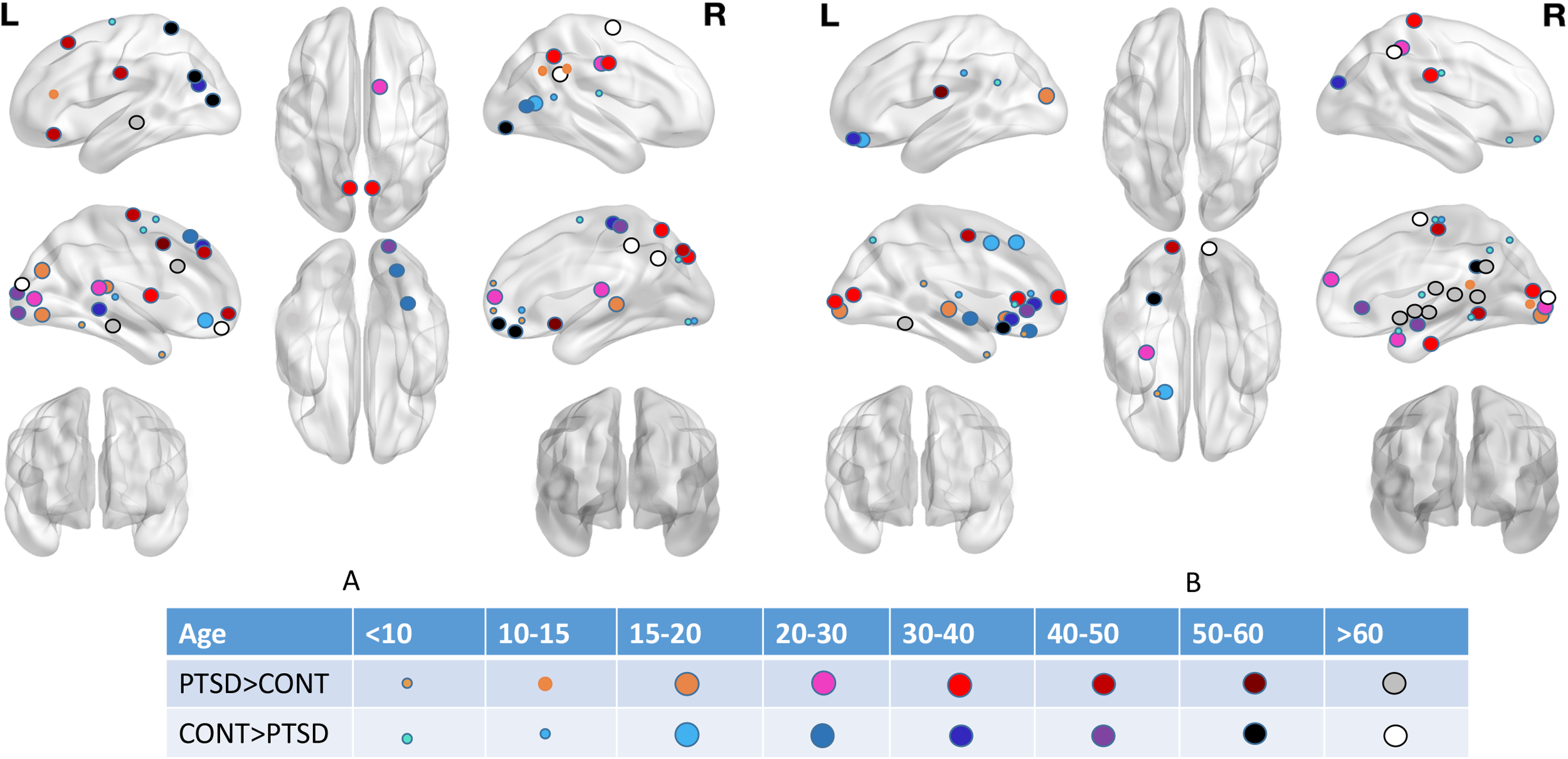
shows a SCN comparison between PTSD subjects and healthy controls across different age groups. A is for cortical thickness (CT) and B is for surface area (SA) measures. Various age groups that are compared include <10 years, 10-15 years, 15-20 years, 20-30 years, 30-40 years, 40-50 years, 50-60 years and > 60 years. Each age group has a different color code as shown in the box below the figure. Views shown at either ends of A and B in the top (first) row depict medial views. Views shown at either ends of A and B in the middle (second) row depict lateral views.

Regions with significant interactions of diagnoses and sex are listed in **Table 5. Table 6** shows the main effect of sex (males versus females regardless of diagnoses); **tables 7 and 8** show how males and females compare in PTSD subjects and control subjects respectively. **Tables 9** and **10** shows results comparing centrality of PTSD and control groups in males and females respectively. We show all these comparisons in **Figure 4**.

**Figure 4.**
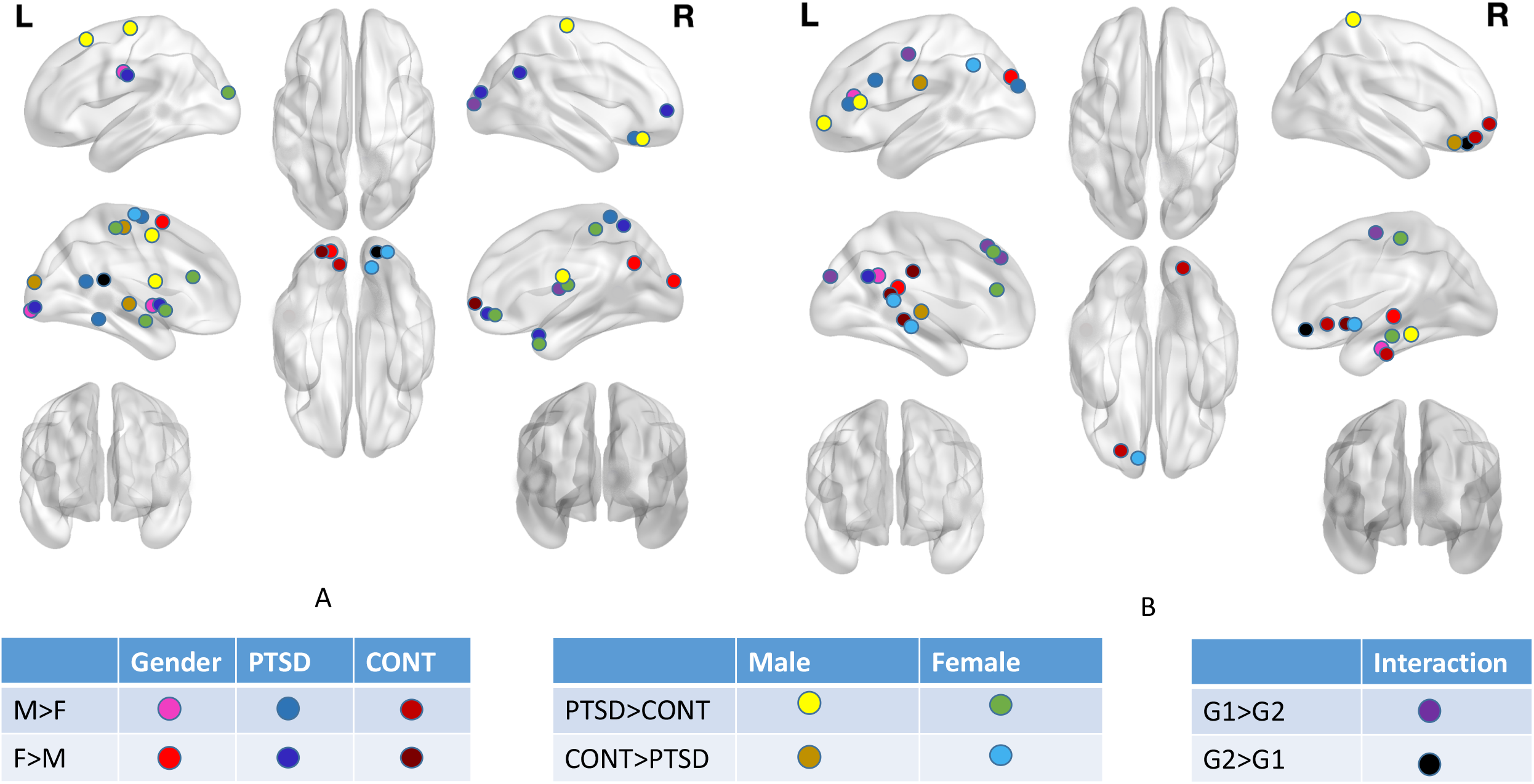
shows diagnoses sex interaction comparisons between PTSD subjects and healthy controls. A is for cortical thickness and B is for surface area measures. Comparisons highlighted in the figure include the main effect of sex, male versus female subjects with PTSD, male versus female healthy controls, PTSD subjects versus healthy controls among males, PTSD subjects versus healthy controls amongst females and then the interaction effect. For the interaction effect, G1 stands for (male and PTSD or female and no PTSD) and G2 stands for (female & PTSD OR male & no PTSD).

Nodes that showed a significant interaction of PTSD and MDD are listed in **Table 11**, while the main effect of comorbid MDD is provided in **Table 12. Table 13** compares PTSD with comorbid MDD to PTSD alone, and **Table 14** compares PTSD with comorbid MDD to controls.

#### 3.2 SA SCN Results

#### 3.2.1 Graph Centrality Based on Pearson’s Correlation Coefficients

Compared to control subjects, PTSD subjects showed higher centrality in the left superior temporal sulcus and middle posterior part of the right cingulate gyrus and sulcus. Compared to PTSD subjects, controls showed higher centrality in the left pericallosal sulcus (**Figure 2 and Table 15**).

**Supplemental Table 13** provides results comparing centrality measures between PTSD and control subjects using negative correlation coefficients as connections.

#### 3.2.2 Interactions

Brain regions with significantly different centrality between PTSD subjects and controls are listed in **Figure 3** and **Supplemental Tables 14** to **21** for age groups <10, 10-15, 15-20, 20-30, 30-40, 40-50, 50-60, and >60 respectively.

Nodes showing a significant interaction between diagnoses and sex are listed in **Table 16. Table 17** shows main effect of sex. **Tables 18** and **19** show comparisons between males and females in PTSD and control groups respectively. **Tables 20** and **21** show comparisons between PTSD and control groups in males and females respectively. We show all of these results in **Figure 4** as well.

Nodes with significant centrality interactions of PTSD diagnoses and MDD diagnosis are listed in **Table 22. Table 23** shows the main effect of MDD (i.e. MDD with and without PTSD vs. no MDD with and without PTSD). **Tables 24** and **25** show how subjects with PTSD and comorbid MDD compare to those with only PTSD as well as how subjects with PTSD and comorbid MDD compare to controls.

## 4. DISCUSSION

To the best of our knowledge, our study is larger than prior SCN studies of PTSD by an order of magnitude and is the first cross-sectional study to examine SCN interactions between PTSD diagnosis and age (**Figure 3**), diagnosis and sex (**Figure 4**), and diagnosis and MDD. Although CT and SA revealed contrasting results (**Figure 2**), both measures pointed to frontal, temporal and occipital involvement, which falls in line with results from previous structural neuroimaging studies in PTSD (Akiki et al., 2017; Fenster, Lebois, Ressler, & Suh, 2018). Supplementary Tables 22 and 23 display regions with overlapping results across age bins in the present study, compared with results from previous studies.

In our analyses, positive correlations with cortical thickness showed higher centrality in left inferior frontal gyrus and left fusiform gyrus, both of which are areas that mediate fear conditioning (Fenster et al., 2018; Morey et al., 2015). This analysis also showed centrality measure differences in anterior occipital sulcus and posterior transverse collateral sulcus. There is sparse literature on how occipital brain regions are affected in PTSD, mainly in the context of visual processing (Neumeister et al., 2017) and flashbacks(Whalley et al., 2013). CT-based SCN showed PTSD to be associated with higher centrality of the right inferior temporal gyrus and left superior temporal gyrus. These regions form a part of the default network (DMN) dorsomedial subsystem (Andrews-Hanna, Smallwood, & Spreng, 2014). With SA, the left superior temporal sulcus (parts of DMN dorsomedial subsystem (Andrews-Hanna et al., 2014)) and right cingulate gyrus and sulcus (part of salience network) showed increased centrality.

Nodes with significantly higher centrality measures represent focal points of communication and could be considered overall drivers of brain activity due to their structural and/or functional connectivity (Lerch et al., 2006; Mechelli, Friston, Frackowiak, & Price, 2005). Since our results show only positive correlations between involved regions, discerning whether all the relevant regions are increasing or decreasing in CT/SA would need further investigation (Zuo et al., 2012). For functional connectivity data, centrality measures offer insights into a given node’s relationship to the entire functional connectome unlike other approaches like seed-based connectivity or independent component analysis(Zuo et al., 2012). Centrality derived from cortical measures comports with white matter tracts(Gong et al., 2012) as well as with gene expression (Romero-Garcia et al., 2018), but such converging evidence is not available for other graph theory measures like efficiency.

In a study that used SCN to investigate 300 children with normal development trajectories who were into four age groups (Zielinski, Gennatas, Zhou, & Seeley, 2010), salience and executive control networks matured earlier in females across all age groups, these networks were less distributed in the first two age groups compared to the latter two(Zielinski et al., 2010). At baseline, females have more pronounced amygdala activation and connectivity to prefrontal cortex compared to males (Helpman et al., 2017). Increased connectivity in the salience network is seen in males with PTSD, with reversal of this pattern seen in females with PTSD (Helpman et al., 2017). In our study, males with PTSD showed higher CT-based centrality than females with PTSD in bilateral central sulci, left temporal lobe, and right straight gyrus (part of vmPFC) as well as higher SA-based centrality in left anterior cingulate and left superior occipital gyrus. Males with PTSD showed higher CT-based centrality than male controls mainly in bilateral frontal lobes. By contrast, female subjects with PTSD showed higher centrality than female controls in bilateral frontal and temporal lobes, including the bilateral insula. Areas like the vmPFC (Kuhn & Gallinat, 2013), anterior cingulate (Clausen et al., 2020) and superior occipital gyrus (Crombie, Ross, Letkiewicz, Sartin-Tarm, & Cisler, 2021) have been implicated in pathophysiology of PTSD. However, they have not shown sex differences in previous studies but showed differences in our SCN analyses. This could be explained by the relatively sparse literature on sex differences in PTSD. Thus, linking sex-specific structural findings of the vmPFC, anterior cingulate, and superior occipital gyrus to PTSD represents a unique contribution of the present study to the literature.

To the best of our knowledge, there are no studies that have investigated brain changes in PTSD across the lifespan. Available evidence in pediatric PTSD points to aberrant activity in hippocampus, amygdala and vmPFC(Herringa, 2017). In our analyses with CT, we found children (<10 years) and adolescents (10-15 years) to have centrality differences in frontal, temporal and parietal nodes. Starting with emerging adults (15-20 years and 20-30 years) and other age groups, we saw multiple nodes having centrality differences in all brain lobes with heavy conglomeration in the occipital regions. This advocates for disrupted frontolimbic circuitry in pediatric PTSD, consistent with previous studies (Herringa, 2017) and disproportionately heavy occipital involvement in adult PTSD (Chao, Lenoci, & Neylan, 2012; Crombie et al., 2021).

Despite high comorbidity rate of 48% (Flory & Yehuda, 2015; Kessler, Sonnega, Bromet, Hughes, & Nelson, 1995), there has also been scant literature investigating impact of comorbid MDD on brain changes in PSTD. Comorbid PTSD and MDD showed higher CT-based centrality in the right anterior cingulate gyrus, right parahippocampal gyrus, left sulcus intermedius primus, and right anterior transverse temporal gyrus; corresponding differences were not identified for SA-based centrality. However, the left insular gyrus was found to have lower SA centrality in subjects with PTSD and comorbid MDD compared to subjects with only PTSD. Our present results show left insula and right anterior cingulate regions having centrality differences between comorbid PTSD and MDD versus PTSD alone. This result is consistent with a study that compared network resting state connectivity between 27 subjects with PTSD and MDD and 23 subjects with only PTSD. In this study, connectivity between the subgenual ACC and perigenual parts of the ACC was increased in PTSD subjects with comorbid MDD compared to those without. Those with comorbid MDD group showed reduced functional connectivity between insula and hippocampus compared to the those without (Kennis, Rademaker, van Rooij, Kahn, & Geuze, 2013).

## Strengths and Limitations

This study has several strengths including a large sample size and multiple cohorts representing diverse trauma types, race, geography, demography, and chronicity by virtue of an extensive network of consortium contributors. However, this strength also represents an inherent weakness in so far as each site had different recruitment practices, different methods for clinical assessment even when using the same instruments was used, and differing inclusion/exclusion criteria. Another strength was our strategy for addressing scanner-specific effects on cortical thickness and surface area with *ComBat* harmonization. *ComBat* applies empirical Bayes to improve the estimation of site parameters and has been shown to effectively remove unwanted sources of scanner and/or site variability (Fortin et al., 2018). We entered PTSD diagnosis, age, and sex as “biological variables” during ComBat harmonization to preserve their associated variability while removing variability associated with site and scanner. Among other limitations, firstly, we were not able to examine thickness in subcortical brain regions. Second, the present method does not allow for examining the effects of individual differences (e.g. PTSD symptom severity) on network characteristics; only a single network and associated network features are available for a particular group (e.g. PTSD males). Inter-regional correlations in network analyses resting-state functional magnetic resonance imaging, provide connectivity information for each subject, whereas SCN inter-regional relationships are available only at the group-level (Sun, Peverill, et al., 2018). Third, our study utilized a cross-sectional design, which limits inferences about the causal relationships between childhood or adult trauma exposure, PTSD status and severity, and CT-based SCN characteristics. Fourth, we did not include the effects of medication, due to inconsistent methods of collecting this information at many sites.

## Conclusions

We compared SCNs using CT and SA between PTSD subjects and controls, including interactions with age, sex and comorbid MDD. Compared to previous studies with sample sizes (~300), we had a large sample, and our study is the first to look at SCN interactions. We demonstrated altered centrality at areas forming parts of dorsomedial DMN and salience networks. We found sex-based centrality differences in left anterior cingulate, left superior occipital cortex and right vmPFC and this represents a unique contribution of our study. We also found comorbid MDD based centrality differences in right anterior cingulate and left insular gyrus. Nodes with significantly altered centrality measures could be translational targets for precision neuromodulation in PTSD, based on sex, illness severity and comorbid MDD if present.

## Data Availability Statement

Data available on request from the authors

## Notes

Conflicts of interest and disclosures PMT received partial research support from Biogen, Inc. (Boston, USA) for research unrelated to the topic of this manuscript. NJ received partial research support from Biogen, Inc. (Boston, USA) for research unrelated to the content of this manuscript. CS is consultant for Boehringer Ingelheim International GmbH. DJS has received research grants and/or consultancy honoraria from Lundbeck and Servier. RJD is the founder and president of, and serves on the board of directors for, the non-profit organization Healthy Minds Innovations, Inc. CGA has served as a consultant, speaker and/or on advisory boards for FSV7, Lundbeck, Psilocybin Labs, Guidepoint, Genentech and Janssen, and editor of Chronic Stress for Sage Publications, Inc.; he has filed a patent for using mTOR inhibitors to augment the effects of antidepressants (filed on August 20, 2018). JHK is a consultant for AbbVie, Inc., Amgen, Astellas Pharma Global Development, Inc., AstraZeneca Pharmaceuticals, Biomedisyn Corporation, Bristol-Myers Squibb, Eli Lilly and Company, Euthymics Bioscience, Inc., Neurovance, Inc., FORUM Pharmaceuticals, Janssen Research & Development, Lundbeck Research USA, Novartis Pharma AG, Otsuka America Pharmaceutical, Inc., Sage Therapeutics, Inc., Sunovion Pharmaceuticals, Inc., and Takeda Industries; is on the Scientific Advisory Board for Lohocla Research Corporation, Mnemosyne Pharmaceuticals, Inc., Naurex, Inc., and Pfizer; is a stockholder in Biohaven Pharmaceuticals; holds stock options in Mnemosyne Pharmaceuticals, Inc.; holds patents for Dopamine and Noradrenergic Reuptake Inhibitors in Treatment of Schizophrenia, US Patent No. 5,447,948 (issued September 5, 1995), and Glutamate Modulating Agents in the Treatment of Mental Disorders, U.S. Patent No. 8,778,979 (issued July 15, 2014); and filed a patent for Intranasal Administration of Ketamine to Treat Depression. U.S. Application No. 14/197,767 (filed on March 5, 2014); US application or Patent Cooperation Treaty international application No. 14/306,382 (filed on June 17, 2014). Filed a patent for using mTOR inhibitors to augment the effects of antidepressants (filed on August 20, 2018)

### Competing Interest Statement

Conflicts of interest and disclosures 
PMT received partial research support from Biogen, Inc. (Boston, USA) for research unrelated to the topic of this manuscript. 
NJ received partial research support from Biogen, Inc. (Boston, USA) for research unrelated to the content of this manuscript. 
CS is consultant for Boehringer Ingelheim International GmbH. DJS has received research grants and/or consultancy honoraria from Lundbeck and Servier. 
RJD is the founder and president of, and serves on the board of directors for, the non-profit organization Healthy Minds Innovations, Inc. 
CGA has served as a consultant, speaker and/or on advisory boards for FSV7, Lundbeck, Psilocybin Labs, Guidepoint, Genentech and Janssen, and editor of Chronic Stress for Sage Publications, Inc.; he has filed a patent for using mTOR inhibitors to augment the effects of antidepressants (filed on August 20, 2018). 
JHK is a consultant for AbbVie, Inc., Amgen, Astellas Pharma Global Development, Inc., AstraZeneca Pharmaceuticals, Biomedisyn Corporation, Bristol-Myers Squibb, Eli Lilly and Company, Euthymics Bioscience, Inc., Neurovance, Inc., FORUM Pharmaceuticals, Janssen Research & Development, Lundbeck Research USA, Novartis Pharma AG, Otsuka America Pharmaceutical, Inc., Sage Therapeutics, Inc., Sunovion Pharmaceuticals, Inc., and Takeda Industries; is on the Scientific Advisory Board for Lohocla Research Corporation, Mnemosyne Pharmaceuticals, Inc., Naurex, Inc., and Pfizer; is a stockholder in Biohaven Pharmaceuticals; holds stock options in Mnemosyne Pharmaceuticals, Inc.; holds patents for Dopamine and Noradrenergic Reuptake Inhibitors in Treatment of Schizophrenia, US Patent No. 5,447,948 (issued September 5, 1995), and Glutamate Modulating Agents in the Treatment of Mental Disorders, U.S. Patent No. 8,778,979 (issued July 15, 2014); and filed a patent for Intranasal Administration of Ketamine to Treat Depression. U.S. Application No. 14/197,767 (filed on March 5, 2014); US application or Patent Cooperation Treaty international application No. 14/306,382 (filed on June 17, 2014). Filed a patent for using mTOR inhibitors to augment the effects of antidepressants (filed on August 20, 2018)

### Summary of Updates

Name of co-author updated from Scott Sponheim to Scott R. Sponheim

## References

Akiki, T. J., Averill, C. L., & Abdallah, C. G. (2017). A Network-Based Neurobiological Model of PTSD: Evidence From Structural and Functional Neuroimaging Studies. Curr Psychiatry Rep, 19(11), 81. doi:10.1007/s11920-017-0840-4

Andrews-Hanna, J. R., Smallwood, J., & Spreng, R. N. (2014). The default network and self-generated thought: component processes, dynamic control, and clinical relevance. Ann N Y Acad Sci, 1316, 29–52. doi:10.1111/nyas.12360

Bassett, D. S., Bullmore, E., Verchinski, B. A., Mattay, V. S., Weinberger, D. R., & Meyer-Lindenberg, A. (2008). Hierarchical organization of human cortical networks in health and schizophrenia. J Neurosci, 28(37), 9239–9248. doi:10.1523/JNEUROSCI.1929-08.2008

Bromis, K., Calem, M., Reinders, A., Williams, S. C. R., & Kempton, M. J. (2018). Meta-Analysis of 89 Structural MRI Studies in Posttraumatic Stress Disorder and Comparison With Major Depressive Disorder. Am J Psychiatry, 175(10), 989–998. doi:10.1176/appi.ajp.2018.17111199

Chao, L. L., Lenoci, M., & Neylan, T. C. (2012). Effects of post-traumatic stress disorder on occipital lobe function and structure. Neuroreport, 23(7), 412–419. doi:10.1097/WNR.0b013e328352025e

Clausen, A. N., Clarke, E., Phillips, R. D., Haswell, C., Workgroup, V. A. M.-A. M., & Morey, R. A. (2020). Combat exposure, posttraumatic stress disorder, and head injuries differentially relate to alterations in cortical thickness in military Veterans. Neuropsychopharmacology, 45(3), 491–498. doi:10.1038/s41386-019-0539-9

Crombie, K. M., Ross, M. C., Letkiewicz, A. M., Sartin-Tarm, A., & Cisler, J. M. (2021). Differential relationships of PTSD symptom clusters with cortical thickness and grey matter volumes among women with PTSD. Sci Rep, 11(1), 1825. doi:10.1038/s41598-020-80776-2

Destrieux, C., Fischl, B., Dale, A., & Halgren, E. (2010). Automatic parcellation of human cortical gyri and sulci using standard anatomical nomenclature. Neuroimage, 53(1), 1–15. doi:10.1016/j.neuroimage.2010.06.010

Fenster, R. J., Lebois, L. A. M., Ressler, K. J., & Suh, J. (2018). Brain circuit dysfunction in post-traumatic stress disorder: from mouse to man. Nat Rev Neurosci, 19(9), 535–551. doi:10.1038/s41583-018-0039-7

Flory, J. D., & Yehuda, R. (2015). Comorbidity between post-traumatic stress disorder and major depressive disorder: alternative explanations and treatment considerations. Dialogues Clin Neurosci, 17(2), 141–150.

Fortin, J. P., Cullen, N., Sheline, Y. I., Taylor, W. D., Aselcioglu, I., Cook, P. A., Shinohara, R. T. (2018). Harmonization of cortical thickness measurements across scanners and sites. Neuroimage, 167, 104–120. doi:10.1016/j.neuroimage.2017.11.024

Gong, G., He, Y., Chen, Z. J., & Evans, A. C. (2012). Convergence and divergence of thickness correlations with diffusion connections across the human cerebral cortex. Neuroimage, 59(2), 1239–1248. doi:10.1016/j.neuroimage.2011.08.017

Grasby, K. L., Jahanshad, N., Painter, J. N., Colodro-Conde, L., Bralten, J., Hibar, D. P., Enhancing NeuroImaging Genetics through Meta-Analysis Consortium Genetics working, g. (2020). The genetic architecture of the human cerebral cortex. Science, 367(6484). doi:10.1126/science.aay6690

Harnett NG, S. J., Fani N, van Rooij SJH, Ely TD, Michopoulos V, Hudak L, Rothbaum AO, Hinrichs R, Winters SJ, Jovanovic T, Rothbaum BO, Nickerson LD, Ressler KJ. (Available online 7 August 2020). Acute posttraumatic symptoms are associated with multimodal neuroimaging structural covariance patterns: a possible role for the neural substrates of visual processing in PTSD. Biological Psychiatry: Cognitive Neuroscience and Neuroimaging, In press, Journal Pre-proof.

He, Y., Chen, Z. J., & Evans, A. C. (2007). Small-world anatomical networks in the human brain revealed by cortical thickness from MRI. Cereb Cortex, 17(10), 2407–2419. doi:10.1093/cercor/bhl149

He, Y., & Evans, A. (2010). Graph theoretical modeling of brain connectivity. Curr Opin Neurol, 23(4), 341–350. doi:10.1097/WCO.0b013e32833aa567

Helpman, L., Zhu, X., Suarez-Jimenez, B., Lazarov, A., Monk, C., & Neria, Y. (2017). Sex Differences in Trauma-Related Psychopathology: a Critical Review of Neuroimaging Literature (2014-2017). Curr Psychiatry Rep, 19(12), 104. doi:10.1007/s11920-017-0854-y

Herringa, R. J. (2017). Trauma, PTSD, and the Developing Brain. Curr Psychiatry Rep, 19(10), 69. doi:10.1007/s11920-017-0825-3

Hughes, K. C., & Shin, L. M. (2011). Functional neuroimaging studies of post-traumatic stress disorder. Expert Rev Neurother, 11(2), 275–285. doi:10.1586/ern.10.198

Kennis, M., Rademaker, A. R., van Rooij, S. J., Kahn, R. S., & Geuze, E. (2013). Altered functional connectivity in posttraumatic stress disorder with versus without comorbid major depressive disorder: a resting state fMRI study. F1000Res, 2, 289. doi:10.12688/f1000research.2-289.v2

Kessler, R. C., Berglund, P., Demler, O., Jin, R., Merikangas, K. R., & Walters, E. E. (2005). Lifetime prevalence and age-of-onset distributions of DSM-IV disorders in the National Comorbidity Survey Replication. Arch Gen Psychiatry, 62(6), 593–602. doi:10.1001/archpsyc.62.6.593

Kessler, R. C., Sonnega, A., Bromet, E., Hughes, M., & Nelson, C. B. (1995). Posttraumatic stress disorder in the National Comorbidity Survey. Arch Gen Psychiatry, 52(12), 1048–1060. doi:10.1001/archpsyc.1995.03950240066012

Koch, S. B., van Zuiden, M., Nawijn, L., Frijling, J. L., Veltman, D. J., & Olff, M. (2016). Aberrant Resting-State Brain Activity in Posttraumatic Stress Disorder: A Meta-Analysis and Systematic Review. Depress Anxiety, 33(7), 592–605. doi:10.1002/da.22478

Konietschke, F., Schwab, K., & Pauly, M. (2020). Small sample sizes: A big data problem in high-dimensional data analysis. Stat Methods Med Res, 962280220970228. doi:10.1177/0962280220970228

Kuhn, S., & Gallinat, J. (2013). Gray matter correlates of posttraumatic stress disorder: a quantitative meta-analysis. Biol Psychiatry, 73(1), 70–74. doi:10.1016/j.biopsych.2012.06.029

Lerch, J. P., Worsley, K., Shaw, W. P., Greenstein, D. K., Lenroot, R. K., Giedd, J., & Evans, A. C. (2006). Mapping anatomical correlations across cerebral cortex (MACACC) using cortical thickness from MRI. Neuroimage, 31(3), 993–1003. doi:10.1016/j.neuroimage.2006.01.042

Liu, H., Petukhova, M. V., Sampson, N. A., Aguilar-Gaxiola, S., Alonso, J., Andrade, L. H., World Health Organization World Mental Health Survey, C. (2017). Association of DSM-IV Posttraumatic Stress Disorder With Traumatic Experience Type and History in the World Health Organization World Mental Health Surveys. JAMA Psychiatry, 74(3), 270–281. doi:10.1001/jamapsychiatry.2016.3783

Logue, M. W., van Rooij, S. J. H., Dennis, E. L., Davis, S. L., Hayes, J. P., Stevens, J. S., Morey, R. A. (2018). Smaller Hippocampal Volume in Posttraumatic Stress Disorder: A Multisite ENIGMA-PGC Study: Subcortical Volumetry Results From Posttraumatic Stress Disorder Consortia. Biol Psychiatry, 83(3), 244–253. doi:10.1016/j.biopsych.2017.09.006

Long, Z., Duan, X., Xie, B., Du, H., Li, R., Xu, Q., Chen, H. (2013). Altered brain structural connectivity in post-traumatic stress disorder: a diffusion tensor imaging tractography study. J Affect Disord, 150(3), 798–806. doi:10.1016/j.jad.2013.03.004

Mechelli, A., Friston, K. J., Frackowiak, R. S., & Price, C. J. (2005). Structural covariance in the human cortex. J Neurosci, 25(36), 8303–8310. doi:10.1523/JNEUROSCI.0357-05.2005

Meng, L., Jiang, J., Jin, C., Liu, J., Zhao, Y., Wang, W., Gong, Q. (2016). Trauma-specific Grey Matter Alterations in PTSD. Sci Rep, 6, 33748. doi:10.1038/srep33748

Morey, R. A., Dunsmoor, J. E., Haswell, C. C., Brown, V. M., Vora, A., Weiner, J., LaBar, K. S. (2015). Fear learning circuitry is biased toward generalization of fear associations in posttraumatic stress disorder. Transl Psychiatry, 5, e700. doi:10.1038/tp.2015.196

Mueller, S. G., Ng, P., Neylan, T., Mackin, S., Wolkowitz, O., Mellon, S., Weiner, M. W. (2015). Evidence for disrupted gray matter structural connectivity in posttraumatic stress disorder. Psychiatry Res, 234(2), 194–201. doi:10.1016/j.pscychresns.2015.09.006

Neumeister, P., Feldker, K., Heitmann, C. Y., Helmich, R., Gathmann, B., Becker, M. P. I., & Straube, T. (2017). Interpersonal violence in posttraumatic women: brain networks triggered by trauma-related pictures. Soc Cogn Affect Neurosci, 12(4), 555–568. doi:10.1093/scan/nsw165

Nievergelt, C. M., Maihofer, A. X., Klengel, T., Atkinson, E. G., Chen, C. Y., Choi, K. W., Koenen, K. C. (2019). International meta-analysis of PTSD genome-wide association studies identifies sex-and ancestry-specific genetic risk loci. Nat Commun, 10(1), 4558. doi:10.1038/s41467-019-12576-w

Philip, N. S., Carpenter, S. L., & Sweet, L. H. (2014). Developing neuroimaging phenotypes of the default mode network in PTSD: integrating the resting state, working memory, and structural connectivity. J Vis Exp(89). doi:10.3791/51651

Romero-Garcia, R., Whitaker, K. J., Vasa, F., Seidlitz, J., Shinn, M., Fonagy, P., Vertes, P. E. (2018). Structural covariance networks are coupled to expression of genes enriched in supragranular layers of the human cortex. Neuroimage, 171, 256–267. doi:10.1016/j.neuroimage.2017.12.060

Rubinov, M., & Sporns, O. (2010a). Complex network measures of brain connectivity: Uses and interpretations. Neuroimage, 52(3), 1059–1069. doi:DOI 10.1016/j.neuroimage.2009.10.003

Rubinov, M., & Sporns, O. (2010b). Complex network measures of brain connectivity: uses and interpretations. Neuroimage, 52(3), 1059–1069. doi:10.1016/j.neuroimage.2009.10.003

Sun, D., Davis, S. L., Haswell, C. C., Swanson, C. A., Mid-Atlantic, M. W., LaBar, K. S., Morey, R. A. (2018). Brain Structural Covariance Network Topology in Remitted Posttraumatic Stress Disorder. Front Psychiatry, 9, 90. doi:10.3389/fpsyt.2018.00090

Sun, D., Haswell, C. C., Morey, R. A., & De Bellis, M. D. (2018). Brain structural covariance network centrality in maltreated youth with PTSD and in maltreated youth resilient to PTSD. Dev Psychopathol, 1–15. doi:10.1017/S0954579418000093

Sun, D., Peverill, M. R., Swanson, C. S., McLaughlin, K. A., & Morey, R. A. (2018). Structural covariance network centrality in maltreated youth with posttraumatic stress disorder. J Psychiatr Res, 98, 70–77. doi:10.1016/j.jpsychires.2017.12.015

Teicher, M. H., Anderson, C. M., Ohashi, K., & Polcari, A. (2014). Childhood maltreatment: altered network centrality of cingulate, precuneus, temporal pole and insula. Biol Psychiatry, 76(4), 297–305. doi:10.1016/j.biopsych.2013.09.016

Tursich, M., Ros, T., Frewen, P. A., Kluetsch, R. C., Calhoun, V. D., & Lanius, R. A. (2015). Distinct intrinsic network connectivity patterns of post-traumatic stress disorder symptom clusters. Acta Psychiatr Scand, 132(1), 29–38. doi:10.1111/acps.12387

Wang, X., Xie, H., Chen, T., Cotton, A. S., Salminen, L. E., Logue, M. W., Liberzon, I. (2020). Cortical volume abnormalities in posttraumatic stress disorder: an ENIGMA-psychiatric genomics consortium PTSD workgroup mega-analysis. Mol Psychiatry. doi:10.1038/s41380-020-00967-1

Whalley, M. G., Kroes, M. C., Huntley, Z., Rugg, M. D., Davis, S. W., & Brewin, C. R. (2013). An fMRI investigation of posttraumatic flashbacks. Brain Cogn, 81(1), 151–159. doi:10.1016/j.bandc.2012.10.002

Yun, J. Y., Boedhoe, P. S. W., Vriend, C., Jahanshad, N., Abe, Y., Ameis, S. H., Kwon, J. S. (2020). Brain structural covariance networks in obsessive-compulsive disorder: a graph analysis from the ENIGMA Consortium. Brain, 143(2), 684–700. doi:10.1093/brain/awaa001

Zielinski, B. A., Gennatas, E. D., Zhou, J., & Seeley, W. W. (2010). Network-level structural covariance in the developing brain. Proc Natl Acad Sci U S A, 107(42), 18191–18196. doi:10.1073/pnas.1003109107

Zuo, X. N., Ehmke, R., Mennes, M., Imperati, D., Castellanos, F. X., Sporns, O., & Milham, M. P. (2012). Network centrality in the human functional connectome. Cereb Cortex, 22(8), 1862–1875. doi:10.1093/cercor/bhr269

