## Supplemental Material for "Structural Covariance Networks in Post-Traumatic Stress Disorder: A Multisite ENIGMA-PGC Study"

**Methods**

**2.2.1 Rating Scale Description**

Most sites used clinician-administered measures such as the Structured Clinical Interview (SCID) or Clinician-Administered PTSD Scale (Weathers et al., 2018; Weathers, Keane, & Davidson, 2001), to ascertain PTSD diagnosis. One site used a psychiatrist diagnosis for PTSD, and a few additional sites used self-report scales such as the PTSD Checklist (PCL; Weathers et al., 2013) and the Davidson Trauma Scale (Davidson et al., 1997). While the majority used DSM-IV criteria, a small subset of sites used DSM-5 criteria (see Supplementary Table S1 for further details). The aforementioned measures reflect PTSD diagnosis and symptoms in the past month before scanning. For depression, sites used a mix of clinician-administered and self-report instruments to diagnose depression. We harmonized depression data by assigning participants to major depressive disorder (MDD) or control groups based on a standardized depression severity cut-off. The majority of sites reported depression severity using the Beck Depression Inventory-II (Beck AT, 1996)(BDI-II), while other commonly used scales included the Center for Epidemiologic Studies Depression Scale (Carleton et al., 2013)(CES-D), Hamilton Depression Rating Scale (Hamilton, 1960), Depression Anxiety Stress Scales (Lovibond & Lovibond, 1995)(DASS), and Children's Depression Inventory(Kovacs, 1985) (CDI). Cutoffs were chosen based on standard cutoffs reported in previously published studies.

**2.2.4 Harmonizing data across sites.**

ComBat was utilized to harmonize CT and SA values by removing scanner effects across study sites while preserving inherent biological associations in the data (Fortin et al., 2018). ComBat achieves harmonization by first modeling expected imaging features as linear combinations of the biological variables and site effects whose error term is further modulated by site-specific scaling factors. Secondly, ComBat applies empirical Bayes to improve the estimation of site parameters for small samples. This method has been shown to effectively remove unwanted sources of scanner and/or site variability while simultaneously increasing the power and reproducibility of subsequent statistical analyses of multi-site MRI studies (Fortin et al., 2018). PTSD diagnosis, age, and sex were designated as biological variables here to preserve the biological variability while removing variability associated with site and scanner. PTSD severity and depression diagnosis were not designated as biological variables as they were highly correlated with PTSD diagnosis, and some participants were missing information on PTSD severity and depression.

**2.2.5 Adjusting for confounding factors**

Subsequently, age, age^2^, sex, site and mean whole-brain CT or SA estimates were regressed from the CT and SA estimates with a linear model (He, Chen, & Evans, 2007). The age^2^ term was designed to adjust for possible nonlinear effects of age on CT or SA. The mean whole-brain CT or SA estimates were included as regressors to minimize the possibility that subjects with a globally higher CT or SA estimate would be more likely to exhibit larger regional CT or SA estimates. Age was not included as a regressor when we investigated the interaction effect between age and PTSD diagnosis. Similarly, the variable sex was not included as a regressor when we investigated the interaction effect between sex and PTSD diagnosis. The residuals of the linear model were entered into the following analyses.

**2.2.6 Centrality Measures**

(a) *Degree centrality* is the number of directly connected nodes, and reflects how much a node serves as a focal point of communication; (b) *betweenness centrality* is the frequency with which a node falls between the shortest connecting path between two other nodes, and reflects the potential of a node to control communication; (c) *closeness centrality*, which was adapted from the distance function in BCT, reflects the normalized number of steps required to access every other node from a given node in a network, and reflects speed with which a node can spread information throughout the network; (d) *eigenvector centrality* provides the node's overall influence, such that the importance of a node is recursively related to the importance of the nodes associated with it.

**2.2.7 Statistical Analyses**

Analyses were performed with MATLAB scripts that we reported previously (Sun, Haswell, Morey, & De Bellis, 2018; Sun, Peverill, Swanson, McLaughlin, & Morey, 2018). We did not compare the four-centrality measures of each node, given that they are highly correlated. Rather, node centrality was considered to differ between groups only if permutation testing-derived p-values were ≤ 0.05 for at least three out of four centrality measures. Our method is similar to, but far more stringent than, methods for controlling Type 1 error used in earlier reports (Teicher, Anderson, Ohashi, & Polcari, 2014). Given that the SCN approach only delineates a single network per participant group, we assessed the reliability (99% confidence interval) of between-group comparisons using jackknife resampling method that is considered optimal according to Teicher et al (Teicher et al., 2014).

Permutation testing was employed to assess significant between-group differences in centrality measures. The method measures the probability that the between-group difference of nodal centrality could have occurred by chance based on 10,000 network comparisons derived by randomly assigning subjects to two groups. We achieved interaction testing with permutation testing with some modifications of the procedure that is used to test main effects. We randomly assigned to one of four groups when investigating interactions between diagnosis and age, diagnosis and sex, or diagnosis and major depression (MDD). In SCN interaction analyses, to calculate the main effect of sex, (1) we split subjects into male and female, calculated the centrality measure differences between groups, (2) shuffled the group labels and calculated new between-group differences and repeated the procedure 1000 times. The p-values are determined by the comparisons between steps (1) and (2). To calculate PTSD x sex interaction, we (1) split the subjects into 4 groups: male & PTSD, male & non-PTSD, female & PTSD, and female & non-PTSD, and calculated the between-group differences, i.e. (PTSD - non-PTSD) male - (PTSD - non-PTSD) female, (2) shuffled the group labels and calculated new between-group differences 1000 times. We compared steps (1) and (2) to get p-values. In a similar manner, we investigated between-group differences of the contrast between the two levels of age (old versus young) or MDD (MDD versus non-MDD) and performed post-hoc analyses on the nodes showing a significant interaction effect.

Supplemental Table 2 - Age by sex and diagnostic group at each study site

| Study Site | PTSD (Mean) | PTSD (SD) | Controls (Mean) | Controls (SD) | Males  (Mean) | Males (SD) | Females (Mean) | Females (SD) |
| --- | --- | --- | --- | --- | --- | --- | --- | --- |
| ADNI-DoD | 68 | 3.6 | 70 | 5.3 | 69.1 | 4.8 | - | - |
| Booster (AMC) | 40.4 | 9.9 | 39.6 | 10 | 41.7 | 10.2 | 38 | 9.3 |
| Columbia | 36.3 | 9.3 | 35.2 | 10.6 | 40.1 | 9.3 | 33.6 | 9.4 |
| Duke University (DeBellis) | 9.9 | 5.3 | 10.5 | 2.6 | 10.2 | 2.8 | 10.5 | 2.4 |
| Minneapolis VAMC | 32 | 7.6 | 33.2 | 8.6 | 33 | 8.3 | 26.9 | 3.2 |
| Duke University | 40.7 | 9.8 | 40 |  | 40 | 10 | 39.6 | 10.2 |
| Ghent | 32.6 | 10.3 | 37.7 | 12.3 | - | - | 37.1 | 12.1 |
| Groningen (Charité Berlin) | 38.2 | 9.7 | - | - | - | - | 38.2 | 9.7 |
| University of Wisconsin (Grupe) | 30.4 | 6.2 | 30.7 | 6.6 | 30.4 | 6.3 | 33.25 | 9 |
| Emory GTP | 37 | 12.3 | 40.8 | 12.2 | 49.2 | 16.7 | 39.1 | 12.1 |
| INTRUST | 38.7 | 10.5 | 34.8 | 13 | 37 | 12.6 | 34.5 | 12.1 |
| University of Wisconsin (Larson) | 28.8 | 8.5 | 35.2 | 11.3 | 32.2 | 10.6 | 34.4 | 11.1 |
| Leiden | 16 | 1.9 | 14.7 | 1.6 | 14.6 | 1.4 | 15.4 | 1.9 |
| Mannheim | 35.9 | 11.8 | - | - | - | - | 35.9 | 11.8 |
| McLean | 38.2 | 12.9 | 35.6 | 10.5 | - | - | 37.5 | 12.3 |
| Muenster | 27.4 | 7 | 26.5 | 7.4 | 28.8 | 8.3 | 26.7 | 7.1 |
| Phan | 31.3 | 9.3 | 34 | 8.9 | 32.6 | 9.1 | - | - |
| McLean (Rosso) | 35.3 | 7.9 | 33.5 | 9.3 | 34.4 | 8.3 | 33.3 | 9.7 |
| University of Toledo | 40.9 | 9.5 | 34.3 | 11.6 | 38.4 | 11.7 | 32.1 | 10.2 |
| UCAS | 51 | 6.7 | 48.2 | 6.8 | 50 | 7.9 | 49.3 | 5.8 |
| University of Cape Town | 30.5 | 7.2 | 28.7 | 6.4 | - | - | 28.9 | 6.4 |
| University of Washington | 13.2 | 2.9 | 14.1 | 3.3 | 13.5 | 3.1 | 14.4 | 2.9 |
| WACO VA | 41 | 11 | 40.7 | 11.6 | 41 | 10.8 | 40 | 13.6 |
| West Haven VA | 34.8 | 9.2 | 34.2 | 9.8 | 34.8 | 9.7 | 32.4 | 7.8 |
| Yale | 31.8 | 7 | 29.4 | 8.2 | 31.1 | 8.1 | 25.5 | 4 |
| UNSW | 39.3 | 11.6 | 40.9 | 13.1 | 43 | 12.2 | 38.8 | 12.8 |
| South Dakota | 28.8 | 7.1 | 29.9 | 6.9 | 30.6 | 6.8 | 23.6 | 4.8 |
| Stellenbosch | 39.4 | 11 | 43.1 | 14.4 | 44.8 | 13.6 | 40 | 12.6 |
| Stanford | 36.8 | 10.3 | - | - | 37.9 | 11.7 | 36.6 | 9.8 |

Supplemental Table 3 – Scanner Details and Acquisition Parameters Across Sites

| Site/Study | Scanner  Manufacturer | Scanner Model | Channel Number in Head Coil | Acquisition Sequence | Voxel size | FOV (mm) | Acquisition Orientation | TR (milliseconds) | TE (milliseconds) | Flip Angle | Slice Thickness | FreeSurfer Version |
| --- | --- | --- | --- | --- | --- | --- | --- | --- | --- | --- | --- | --- |
| Duke University | GE | MR750 3T | 8 | FSPGR BRAVO | 0.9375x0.9375x0.9375 | 256x256 | Axial | 8160 | 3.2 | 12 | 1 | 5.3 |
|  | GE | EXCITE HD 3T | 8 | FSPGR BRAVO | 1 x 1 x 1 | 240x240 | Axial | 8148; 7840; 8160 | 3.2; 2.9; 3.2 * | 12 | 1 | 5.3 |
|  | GE | 4T LX Nvi | 8 | Spin-echo co-planar | 1x1x1.9 | 240x240 | Axial | 12000 | 5.4 | 20 | 2 | 5.3 |
|  | Philips | Ingenia | 8 | 3D TFE SENSE | 0.9375x0.9375x1 | 240 x 240 | Axial | 8148 | 3.7 | 8 | 1 | 5.3 |
| McLean | Siemens | Trio | 12 | MEMPRAGE tfl_mgh_multiecho | 1x1x1 | 256 x 256 | Sagittal | 2530 | 1.64,3.5,5.36,7.22* | 7 | 1 | 5.3 |
| Toledo | GE | SignaX | 8 | SPGR | 1x1x1 mm | 256x256 | Axial | 7900 | 3 | 9 | 1 | 5.3 |
| West Haven | 3T Siemens | Tim Trio | 12 | MPRAGE | 1x1x1 | 256x256 | Sagittal | 2530 | 2.71 | 7 | 1 | 5.3 |
| Yale | Siemens 3T | Tim Trio | 32 | MPRAGE | 1x1x1 | 256x256 | Sagittal | 2500 | 2.77 | 7 | 1 | 5.3 |
| UCAS | Philips | Achieva | 8 | EPI | 1x1x1 | 220X220 | Axial | 8500 | 3.7 | 8 | 2 | 5.3 |
| ADNI-DoD | GE | GE Discovery MR750w | 40 | FSPGR | 1x1x1.2 | 256x256 | Sagittal | 7652 | 3.104 | 11 | 1.2 | 6 |
|  | GE | GE Signa HDxt | 8 | SPGR | 1x1x1.2 | 256x256 | Sagittal | 6984 | 2.848 | 11 | 1.2 | 6 |
|  | Siemens | Siemens TrioTim | 12 | MPRAGE | 1x1x1.2 | 256x255 | Sagittal | 2300 | 2.98 | 9 | 1.2 | 6 |
|  | GE | GE Discovery MR750 | 8 | SPGR | 1x1x1.2 | 256x256 | Sagittal | 7340 | 3.036 | 11 | 1.2 | 6 |
| Emory GTP | Siemens | TIM Trio | 12 | MPRAGE | 1x1x1 | 224x256 | Axial | 2600 | 3 | 8 | 1 | 5.3 |
| Stellenbosch | Siemens | Allegra | 4 | MPRAGE | 1x1x1 | 256x256 | Sagittal | 2530 | 1.5; 3.2; 4.9; 6.6* | 7 | 1 | 5.3 |
|  | Siemens | Skyra | 32 | MPRAGE | 1x1x1 | 280x280 | Sagittal | 2530 | 1.63; 3.47; 5.31; 7.15* | 7 | 1 | 5.3 |
| Columbia | GE | 1.5T SIGNA EXCITE | 8 | SPGR | 3.5X3.5X2.2 | 224X224 | Axial | 3000 | 30 | 84 | 2.2 | 5.3 |
| Ghent | Siemens | TimTrio | 32 | MPRAGE | 1x1x1 | 256 x 256 | Transversal | 2250 | 4.18 | 9 | 1 | 5.3 |
| Univ of Cape Town | Siemens | Skyra | 4 | MPRAGE | 1x1x1.5 | 256x256 | Sagittal | 2530 | 1.69; 3.55; 5.41; 7.27* | 7 | 1.5 | 5.3 |
|  | Siemens | Allegra | 4 | MPRAGE | 1x1x1.5 | 256x256 | Sagittal | 2000 | 1.53; 3.21; 4.89; 6.57* | 20 | 1.5 | 5.3 |
| INTRUST | GE | multiple | 8 | SPGR-BRAVO | 1x1x1 | 256x256 | Sagittal | 9150 | 3.7 | 10 | 1 | 5.3 |
|  | Siemens | multiple | 12 | MPRAGE | 1x1x1 | 256x256 | Sagittal | 2530 | 3.32 | 7 | 1 | 5.3 |
|  | Philips | multiple | 8 | T1W_3D_TFESENSE | 1x1x1 | 256x256 | Sagittal | 7600 | 3.5 | 7 | 1 | 5.3 |
| UNSW | GE | Signa Hdx | 8 | 3D SPGR | 1x1x1 | 256 x 256 | Sagittal | 8300 | 3.24 | 11 | 1 | 5.3 |
| Leiden | Philips | Achieva | 8 | gradient echo T1-weightedEPI | 1.17x1.17x1.2 | 224x177x168 | Sagittal | 9800 | 4.6 | 8 | 1.2 | 5.3 |
| University of Washington | Siemens | Tim Trio | 32 | MEMPRAGE | 1x1x1 | 220x220 | Sagittal | 2530 | 1.6-7 | 7 | 1 | 5.3 |
|  | Philips | Achieva | 32 | MEMPRAGE | 1x1x1 | 256x256 | Sagittal | 2530 | 1.6 - 7/3.5* | 7 | 1 | 5.3 |
| Duke University (DeBellis) | Siemens | Tim Trio | 8 | MEMPRAGE | 1x1x1 | 256x256 | Axial | 1750 | 5.17 | 20 | 1 | 5.3 |
| Mannheim | Siemens | Trio 3T | 32 | SPGR | 1x1x1 | 192x192 | Axial | 2000 | 3 | 80 | 3 | 5.3 |
| University of Wisconsin-Madison | GE | X750 Discovery | 8 | BRAVO | 1x1x1 | 256x256 | Axial | 8200 | 3.18 | 12 | 1 | 5.3 |
| Groningen | Siemens | TrioTim | 12 | MPRAGE | 1x1x1 | 256X256 | Sagittal | 1900 | 2.5 | 9 | 1 | 5.3 |
| Booster (AMC) | Philips | Achieva | 32 | FAST MPRAGE | 1x1x1 | 240x188 | Axial | 8200 | 3.8 | 8 | 1 | 5.3 |
| McLean (Rosso) | Siemens | Tim Trio | 32 | MEMPRAGE | 1.3X1X1.3 | 256X128 | Sagittal | 2530 | 3.31 | 7 | 1.33 | 5.3 |
| Stanford | GE | MR750 3T | 8 | 3D SPGR | 1.5x0.9x1.1 | 220X220; 240x240 | Coronal | 8000; 8600 | 3.6; 3.4* | 15 | 1.1 | 5.3 |
| Minneapolis VA | Siemens | TimTrio | 12 | MPRAGE | 1x1x1 | 256X256 | Coronal | 2530 | 3.7 | 7 | 1 | 6 |
| Muenster | Siemens | Magnetom Prisma | HE1-4 | MPRAGE | 1x1x1 | 256 X 256 | Sagittal | 2130 | 2.28 | 8 | 1 | 5.3 |
| WACO VA | Philips | Achieva | 16 | MPRAGE | 1x1x1 | 288x288 | Sagittal | 2400 | 3.08 | 90 | 1 | 6 |
| Univ. of Wisconsin-Milwaukee | GE | MR750 Signa Excite | 32 | SPGR | 1x0.9375x0.9375 | 240 X240 | Sagittal | 8200 | 3.2 | 12 | 1 | 5.3 |
| Univ. South Dakota | Siemens | Skyra | 20 | MPRAGE | 1x1x1 | 240x240; 256x256 | Sagittal, interleaved | 1900 | 2.13 | 9 | 0.9 | 5.3 |
| University of Illinois-Chicago | GE | Signa | 8 | Gradient-echo spiral pulse | 1x1x1 | 240x240 | Coronal | 2500 | 6.6 | 90 | 1 | 5.3 |

*- multi-echo acquisitions

Supplemental Table 4- Cortical Thickness – Negative Correlations

| PTSD>CONT | | | | | | | | |
| --- | --- | --- | --- | --- | --- | --- | --- | --- |
| Nodes | Deg_P | Deg_C | Bet_P | Bet_C | Clo_P | Clo_C | Eig_P | Eig_C |
| L_G_and_S_subcentral | 15 | 5 | 53.45 | 5.54 | 74.17 | 65.58 | 0.06 | 0.02 |
| L_G_precentral | 27 | 14 | 328.67 | 40.4 | 83.67 | 72.08 | 0.10 | 0.06 |
| L_S_front_middle | 36 | 25 | 445.26 | 229.16 | 89.67 | 81.83 | 0.13 | 0.09 |
| R_G_and_S_paracentral | 23 | 11 | 195.41 | 33.93 | 80.83 | 72.67 | 0.09 | 0.04 |
| R_G_Ins_lg_and_S_cent_ins | 33 | 21 | 351.01 | 163.01 | 86 | 78.33 | 0.12 | 0.07 |
| R_G_occipital_sup | 30 | 21 | 268.79 | 145.31 | 85.17 | 77.33 | 0.12 | 0.09 |
| R_S_front_inf | 29 | 19 | 459.23 | 135.86 | 85 | 77.33 | 0.10 | 0.06 |
| CONT>PTSD | | | | | | | | |
| L_S_temporal_sup | 26 | 38 | 241.22 | 521.53 | 83.67 | 90.5 | 0.09 | 0.14 |
| R_G_and_S_cingul_mid_Ant | 14 | 30 | 37.51 | 199.27 | 71.83 | 84 | 0.06 | 0.12 |
| R_G_front_sup | 23 | 35 | 284.22 | 473.46 | 81 | 88.5 | 0.07 | 0.11 |
| R_G_subcallosal | 3 | 11 | 1.57 | 25.55 | 58.58 | 72.75 | 0.01 | 0.05 |
| R_Pole_occipital | 15 | 24 | 44.84 | 210.53 | 73.92 | 81 | 0.07 | 0.09 |
| R_S_collat_transv_post | 8 | 17 | 42.04 | 138.56 | 67.08 | 76.83 | 0.02 | 0.06 |
| R_S_oc_sup_and_transversal | 14 | 24 | 34.58 | 213.12 | 73 | 81.83 | 0.06 | 0.1 |
| R_S_occipital_ant | 7 | 19 | 24.5 | 110.05 | 66.42 | 77.08 | 0.02 | 0.08 |
| R_S_temporal_sup | 21 | 30 | 137.29 | 264.95 | 80.17 | 85.17 | 0.08 | 0.12 |

| PTSD>CONT | | | |
| --- | --- | --- | --- |
| p_degree | p_betweenness | p_closeness | p_eigenvector |
| 0.01 | 0.05 | 0.07 | 0.01 |
| 0.01 | 0.01 | 0.003 | 0.03 |
| 0.004 | 0.02 | 0.003 | 0.01 |
| 0.02 | 0.02 | 0.06 | 0.04 |
| 0.01 | 0.07 | 0.03 | 0.01 |
| 0.05 | 0.11 | 0.02 | 0.04 |
| 0.04 | 0.01 | 0.04 | 0.05 |
| CONT > PTSD | | | |
| 0.002 | 0.03 | 0.003 | 0.0004 |
| 0.02 | 0.02 | 0.01 | 0.02 |
| 0.03 | 0.2 | 0.04 | 0.02 |
| 0.01 | 0.04 | 0.01 | 0.002 |
| 0.04 | 0.05 | 0.04 | 0.1 |
| 0.04 | 0.13 | 0.03 | 0.01 |
| 0.04 | 0.04 | 0.02 | 0.06 |
| 0.002 | 0.04 | 0.01 | 0.0004 |
| 0.02 | 0.1 | 0.03 | 0.01 |

Deg_P – Degree centrality for PTSD subjects, Deg_C – Degree centrality for control subjects, Bet_P – Betweenness centrality for PTSD subjects, Bet_C – Betweenness centrality for control subjects, Clo_P – Closeness centrality for PTSD subjects, Clo_C – Closeness centrality for control subjects, Eig_P – Eigen vector centrality for PTSD subjects, Eig_C – Eigen vector centrality for control subjects, p_degree – p value for degree centrality comparison, p_betweenness – p value for betweenness centrality comparison, p_closeness – p value for closeness centrality comparison, p_eigenvector – p value for eigenvector centrality comparison.

Supplemental Table 5- Cortical Thickness - Positive Correlations

**[Insert supplemental figure 1 here]**

Age <10 years

| PTSD>CONT | | | | | | | | |
| --- | --- | --- | --- | --- | --- | --- | --- | --- |
| Nodes | Deg_P | Deg_C | Bet_P | Bet_C | Clo_P | Clo_C | Eig_P | Eig_C |
| R_Lat_fis_ant_Horizont | 26.00 | 9.00 | 225.88 | 149.18 | 77.83 | 66.17 | 0.22 | 0.03 |
| R_S_front_middle | 33.00 | 17.00 | 612.14 | 284.70 | 82.08 | 73.83 | 0.23 | 0.07 |
| R_G_and_S_transv_frontopol | 20.00 | 7.00 | 571.83 | 72.76 | 76.17 | 59.75 | 0.10 | 0.01 |
| L_G_temporal_middle | 19.00 | 9.00 | 432.30 | 165.23 | 74.00 | 64.42 | 0.04 | 0.01 |
| L_Pole_temporal | 19.00 | 4.00 | 313.71 | 18.72 | 73.17 | 55.33 | 0.03 | 0.01 |
| CONT > PTSD | | | | | | | | |
| L_S_central | 7.00 | 27.00 | 89.36 | 483.82 | 62.92 | 79.25 | 0.02 | 0.17 |
| R_G_and_S_occipital_inf | 4.00 | 11.00 | 38.75 | 314.70 | 60.42 | 69.75 | 0.01 | 0.02 |
| R_G_pariet_inf_Angular | 13.00 | 23.00 | 79.97 | 290.96 | 66.83 | 77.75 | 0.02 | 0.13 |
| R_G_precentral | 11.00 | 28.00 | 75.72 | 279.58 | 66.75 | 79.17 | 0.02 | 0.18 |
| R_S_pericallosal | 5.00 | 13.00 | 31.65 | 401.54 | 57.58 | 68.25 | 0.01 | 0.01 |
| L_G_and_S_paracentral | 8.00 | 23.00 | 47.85 | 203.17 | 62.50 | 75.75 | 0.02 | 0.13 |
| L_G_precentral | 12.00 | 31.00 | 108.19 | 478.48 | 69.08 | 82.50 | 0.03 | 0.19 |

| PTSD > CONT | | | |
| --- | --- | --- | --- |
| p_degree | p_betweenness | p_closeness | p_eigenvector |
| 0.02 | 0.29 | 0.05 | 0.002 |
| 0.0008 | 0.01 | 0.02 | 0.002 |
| 0.03 | 0.0008 | 0.01 | 0.03 |
| 0.03 | 0.03 | 0.04 | 0.19 |
| 0.005 | 0.02 | 0.01 | 0.33 |
| CONT > PTSD | | | |
| 0.01 | 0.01 | 0.01 | 0.05 |
| 0.01 | 0.005 | 0.03 | 0.27 |
| 0.07 | 0.04 | 0.03 | 0.04 |
| 0.02 | 0.04 | 0.03 | 0.005 |
| 0.03 | 0.04 | 0.04 | 0.21 |
| 0.01 | 0.07 | 0.01 | 0.04 |
| 0.02 | 0.12 | 0.04 | 0.01 |

Deg_P – Degree centrality for PTSD subjects, Deg_C – Degree centrality for control subjects, Bet_P – Betweenness centrality for PTSD subjects, Bet_C – Betweenness centrality for control subjects, Clo_P – Closeness centrality for PTSD subjects, Clo_C – Closeness centrality for control subjects, Eig_P – Eigen vector centrality for PTSD subjects, Eig_C – Eigen vector centrality for control subjects, p_degree – p value for degree centrality comparison, p_betweenness – p value for betweenness centrality comparison, p_closeness – p value for closeness centrality comparison, p_eigenvector – p value for eigenvector centrality comparison.

Supplemental Table 6- Cortical Thickness -Positive Correlations

10-15 years

| PTSD >CONT | | | | | | | | |
| --- | --- | --- | --- | --- | --- | --- | --- | --- |
| Nodes | Deg_P | Deg_C | Bet_P | Bet_C | Clo_P | Clo_C | Eig_P | Eig_C |
| R_S_subparietal | 25.00 | 11.00 | 543.26 | 137.69 | 80.33 | 70.08 | 0.12 | 0.05 |
| R_G_precuneus | 21.00 | 10.00 | 304.97 | 35.79 | 77.17 | 66.58 | 0.14 | 0.07 |
| L_G_and_S_cingul_Ant | 25.00 | 8.00 | 383.74 | 151.62 | 78.75 | 63.50 | 0.17 | 0.02 |
| CONT > PTSD | | | | | | | | |
| L_G_temp_sup_G_T_transv | 7.00 | 18.00 | 55.44 | 671.16 | 60.75 | 70.83 | 0.01 | 0.03 |
| R_G_and_S_occipital_inf | 8.00 | 23.00 | 60.10 | 348.43 | 66.33 | 76.33 | 0.02 | 0.13 |
| R_G_cingul_Post_ventral | 3.00 | 18.00 | 8.90 | 154.76 | 52.92 | 69.42 | 0.01 | 0.05 |
| R_Lat_fis_ant_Vertical | 6.00 | 8.00 | 57.39 | 97.83 | 63.08 | 66.33 | 0.01 | 0.03 |

| PTSD>CONT | | | |
| --- | --- | --- | --- |
| p_degree | p_betweenness | p_closeness | p_eigenvector |
| 0.03 | 0.01 | 0.04 | 0.17 |
| 0.03 | 0.07 | 0.02 | 0.04 |
| 0.004 | 0.06 | 0.02 | 0.01 |
| CONT > PTSD | | | |
| 0.08 | 0.05 | 0.04 | 0.03 |
| 0.02 | 0.05 | 0.06 | 0.01 |
| 0.003 | 0.10 | 0.002 | 0.02 |
| 0.04 | 0.09 | 0.05 | 0.01 |

Deg_P – Degree centrality for PTSD subjects, Deg_C – Degree centrality for control subjects, Bet_P – Betweenness centrality for PTSD subjects, Bet_C – Betweenness centrality for control subjects, Clo_P – Closeness centrality for PTSD subjects, Clo_C – Closeness centrality for control subjects, Eig_P – Eigen vector centrality for PTSD subjects, Eig_C – Eigen vector centrality for control subjects, p_degree – p value for degree centrality comparison, p_betweenness – p value for betweenness centrality comparison, p_closeness – p value for closeness centrality comparison, p_eigenvector – p value for eigenvector centrality comparison.

Supplemental Table 7- Cortical Thickness - Positive Correlations

15-20 years

| PTSD >CONT | | | | | | | | |
| --- | --- | --- | --- | --- | --- | --- | --- | --- |
| Nodes | Deg_P | Deg_C | Bet_P | Bet_C | Clo_P | Clo_C | Eig_P | Eig_C |
| L_G_and_S_occipital_inf | 23.00 | 13.00 | 333.25 | 111.42 | 76.00 | 66.50 | 0.15 | 0.05 |
| L_G_and_S_subcentral | 20.00 | 8.00 | 264.19 | 131.01 | 76.50 | 63.92 | 0.13 | 0.03 |
| L_G_pariet_inf_Angular | 26.00 | 18.00 | 484.76 | 189.30 | 79.33 | 72.58 | 0.17 | 0.10 |
| R_G_temp_sup_Lateral | 24.00 | 10.00 | 414.19 | 161.06 | 77.83 | 66.58 | 0.13 | 0.03 |
| CONT > PTSD | | | | | | | | |
| R_S_parieto_occipital | 7.00 | 17.00 | 33.47 | 260.39 | 60.42 | 74.17 | 0.03 | 0.11 |
| L_G_front_inf_Orbital | 5.00 | 11.00 | 59.67 | 305.28 | 59.75 | 66.50 | 0.02 | 0.03 |

| PTSD > CONT | | | |
| --- | --- | --- | --- |
| p_degree | p_betweenness | p_closeness | p_eigenvector |
| 0.05 | 0.09 | 0.04 | 0.04 |
| 0.05 | 0.20 | 0.04 | 0.04 |
| 0.05 | 0.02 | 0.04 | 0.10 |
| 0.005 | 0.03 | 0.02 | 0.02 |
| CONT > PTSD | | | |
| 0.01 | 0.01 | 0.002 | 0.004 |
| 0.01 | 0.002 | 0.03 | 0.11 |

Deg_P – Degree centrality for PTSD subjects, Deg_C – Degree centrality for control subjects, Bet_P – Betweenness centrality for PTSD subjects, Bet_C – Betweenness centrality for control subjects, Clo_P – Closeness centrality for PTSD subjects, Clo_C – Closeness centrality for control subjects, Eig_P – Eigen vector centrality for PTSD subjects, Eig_C – Eigen vector centrality for control subjects, p_degree – p value for degree centrality comparison, p_betweenness – p value for betweenness centrality comparison, p_closeness – p value for closeness centrality comparison, p_eigenvector – p value for eigenvector centrality comparison.

Supplemental Table 8- Cortical Thickness - Positive Correlations

20-30 years

| PTSD >CONT | | | | | | | | |
| --- | --- | --- | --- | --- | --- | --- | --- | --- |
| Nodes | Deg_P | Deg_C | Bet_P | Bet_C | Clo_P | Clo_C | Eig_P | Eig_C |
| R_S_orbital_lateral | 20.00 | 12.00 | 374.38 | 28.01 | 73.17 | 57.17 | 0.03 | 0.01 |
| L_G_and_S_subcentral | 21.00 | 7.00 | 607.99 | 105.07 | 75.37 | 56.82 | 0.11 | 0.01 |
| L_G_occipital_middle | 32.00 | 23.00 | 540.90 | 209.60 | 79.50 | 69.95 | 0.21 | 0.17 |
| R_G_and_S_paracentral | 30.00 | 17.00 | 413.05 | 349.07 | 79.20 | 68.58 | 0.21 | 0.11 |
| R_G_and_S_subcentral | 17.00 | 8.00 | 413.86 | 311.78 | 72.37 | 59.62 | 0.07 | 0.01 |
| R_G_and_S_cingul_mid_Post | 17.00 | 9.00 | 318.95 | 191.87 | 72.78 | 61.83 | 0.10 | 0.01 |
| CONT > PTSD | | | | | | | | |
| R_S_oc_temp_lat | 4.00 | 16.00 | 6.54 | 255.52 | 53.57 | 67.58 | 0.01 | 0.07 |
| R_S_parieto_occipital | 18.00 | 29.00 | 87.35 | 554.70 | 68.70 | 75.67 | 0.11 | 0.20 |
| R_S_temporal_inf | 5.00 | 14.00 | 46.41 | 202.86 | 58.17 | 66.00 | 0.01 | 0.03 |
| L_G_front_sup | 15.00 | 26.00 | 143.80 | 1265.10 | 67.12 | 76.17 | 0.04 | 0.05 |

| PTSD > CONT | | | |
| --- | --- | --- | --- |
| p_degree | p_betweenness | p_closeness | p_eigenvector |
| 0.01 | 0.01 | 0.0002 | 0.11 |
| 0.01 | 0.03 | 0.01 | 0.0004 |
| 0.01 | 0.02 | 0.001 | 0.04 |
| 0.03 | 0.32 | 0.02 | 0.03 |
| 0.03 | 0.34 | 0.03 | 0.001 |
| 0.02 | 0.20 | 0.02 | 0.0004 |
| CONT > PTSD | | | |
| 0.001 | 0.02 | 0.003 | 0.003 |
| 0.04 | 0.01 | 0.04 | 0.03 |
| 0.002 | 0.02 | 0.01 | 0.01 |
| 0.02 | 0.002 | 0.01 | 0.35 |

Deg_P – Degree centrality for PTSD subjects, Deg_C – Degree centrality for control subjects, Bet_P – Betweenness centrality for PTSD subjects, Bet_C – Betweenness centrality for control subjects, Clo_P – Closeness centrality for PTSD subjects, Clo_C – Closeness centrality for control subjects, Eig_P – Eigen vector centrality for PTSD subjects, Eig_C – Eigen vector centrality for control subjects, p_degree – p value for degree centrality comparison, p_betweenness – p value for betweenness centrality comparison, p_closeness – p value for closeness centrality comparison, p_eigenvector – p value for eigenvector centrality comparison.

Supplemental Table 9- Cortical Thickness - Positive Correlations

30-40 years

| PTSD > CONT | | | | | | | | |
| --- | --- | --- | --- | --- | --- | --- | --- | --- |
| Nodes | Deg_P | Deg_C | Bet_P | Bet_C | Clo_P | Clo_C | Eig_P | Eig_C |
| R_G_pariet_inf_Angular | 28.00 | 17.00 | 813.68 | 93.36 | 80.50 | 69.00 | 0.18 | 0.12 |
| R_S_intrapariet_and_P_trans | 41.00 | 29.00 | 822.67 | 237.96 | 86.33 | 76.58 | 0.25 | 0.20 |
| R_S_subparietal | 19.00 | 9.00 | 99.05 | 33.01 | 71.92 | 62.00 | 0.14 | 0.06 |
| L_G_front_inf_Opercular | 11.00 | 5.00 | 359.46 | 110.38 | 68.58 | 55.83 | 0.03 | 0.004 |
| L_G_oc_temp_med_Parahip | 12.00 | 4.00 | 140.66 | 39.60 | 64.33 | 50.73 | 0.02 | 0.002 |
| R_G_and_S_paracentral | 18.00 | 9.00 | 221.80 | 20.81 | 72.75 | 61.67 | 0.11 | 0.05 |
| R_G_oc_temp_med_Parahip | 12.00 | 5.00 | 565.42 | 8.00 | 62.50 | 50.68 | 0.01 | 0.003 |
| CONT > PTSD | | | | | | | | |
| R_S_central | 14.00 | 24.00 | 94.64 | 539.37 | 66.92 | 75.58 | 0.08 | 0.15 |
| L_G_cuneus | 17.00 | 25.00 | 78.96 | 162.09 | 65.52 | 72.00 | 0.09 | 0.18 |
| R_S_collat_transv_post | 2.00 | 13.00 | 1.00 | 174.06 | 44.97 | 65.25 | 0.004 | 0.07 |
| L_S_front_sup | 13.00 | 22.00 | 24.78 | 534.83 | 64.33 | 75.42 | 0.04 | 0.06 |
| L_S_temporal_sup | 19.00 | 28.00 | 281.04 | 810.15 | 72.50 | 78.17 | 0.08 | 0.11 |

| PTSD > CONT | | | |
| --- | --- | --- | --- |
| p_degree | p_betweenness | p_closeness | p_eigenvector |
| 0.02 | 0.005 | 0.003 | 0.04 |
| 0.004 | 0.01 | 0.001 | 0.01 |
| 0.01 | 0.38 | 0.01 | 0.005 |
| 0.03 | 0.11 | 0.01 | 0.04 |
| 0.02 | 0.21 | 0.01 | 0.02 |
| 0.02 | 0.04 | 0.01 | 0.03 |
| 0.02 | 0.01 | 0.03 | 0.16 |
| CONT > PTSD | | | |
| 0.04 | 0.09 | 0.04 | 0.04 |
| 0.05 | 0.15 | 0.04 | 0.03 |
| 0.03 | 0.02 | 0.003 | 0.06 |
| 0.03 | 0.03 | 0.01 | 0.31 |
| 0.03 | 0.02 | 0.05 | 0.18 |

Deg_P – Degree centrality for PTSD subjects, Deg_C – Degree centrality for control subjects, Bet_P – Betweenness centrality for PTSD subjects, Bet_C – Betweenness centrality for control subjects, Clo_P – Closeness centrality for PTSD subjects, Clo_C – Closeness centrality for control subjects, Eig_P – Eigen vector centrality for PTSD subjects, Eig_C – Eigen vector centrality for control subjects, p_degree – p value for degree centrality comparison, p_betweenness – p value for betweenness centrality comparison, p_closeness – p value for closeness centrality comparison, p_eigenvector – p value for eigenvector centrality comparison.

Supplemental Table 10- Cortical Thickness - Positive Correlations

40-50 years

| PTSD > CONT | | | | | | | | |
| --- | --- | --- | --- | --- | --- | --- | --- | --- |
| Nodes | Deg_P | Deg_C | Bet_P | Bet_C | Clo_P | Clo_C | Eig_P | Eig_C |
| L_G_and_S_cingul_mid_Post | 23.00 | 11.00 | 812.19 | 211.43 | 77.42 | 66.17 | 0.13 | 0.03 |
| L_G_precentral | 23.00 | 15.00 | 676.36 | 75.22 | 78.50 | 68.50 | 0.13 | 0.08 |
| L_G_rectus | 20.00 | 13.00 | 518.87 | 63.29 | 67.50 | 58.87 | 0.07 | 0.01 |
| L_S_front_sup | 27.00 | 15.00 | 620.61 | 160.27 | 77.83 | 68.42 | 0.12 | 0.07 |
| L_S_orbital_lateral | 17.00 | 7.00 | 118.82 | 41.35 | 66.50 | 55.87 | 0.07 | 0.01 |
| R_G_pariet_inf_Angular | 27.00 | 14.00 | 479.24 | 228.79 | 79.25 | 69.92 | 0.16 | 0.07 |
| CONT > PTSD | | | | | | | | |
| R_S_central | 12.00 | 22.00 | 61.30 | 356.08 | 65.33 | 73.08 | 0.07 | 0.14 |
| R_S_collat_transv_post | 7.00 | 21.00 | 33.41 | 340.99 | 57.50 | 69.25 | 0.02 | 0.10 |
| L_S_oc_middle_and_Lunatus | 12.00 | 25.00 | 40.96 | 293.62 | 63.92 | 73.00 | 0.07 | 0.16 |
| L_S_oc_sup_and_transversal | 16.00 | 34.00 | 55.11 | 631.90 | 69.08 | 80.00 | 0.12 | 0.21 |

| PTSD > CONT | | | |
| --- | --- | --- | --- |
| p_degree | p_betweenness | p_closeness | p_eigenvector |
| 0.002 | 0.01 | 0.01 | 0.0008 |
| 0.04 | 0.02 | 0.01 | 0.11 |
| 0.03 | 0.02 | 0.05 | 0.01 |
| 0.002 | 0.01 | 0.02 | 0.10 |
| 0.03 | 0.21 | 0.03 | 0.01 |
| 0.01 | 0.16 | 0.04 | 0.02 |
| CONT > PTSD | | | |
| 0.05 | 0.03 | 0.03 | 0.09 |
| 0.03 | 0.06 | 0.02 | 0.03 |
| 0.01 | 0.02 | 0.01 | 0.02 |
| 0.001 | 0.01 | 0.002 | 0.01 |

Deg_P – Degree centrality for PTSD subjects, Deg_C – Degree centrality for control subjects, Bet_P – Betweenness centrality for PTSD subjects, Bet_C – Betweenness centrality for control subjects, Clo_P – Closeness centrality for PTSD subjects, Clo_C – Closeness centrality for control subjects, Eig_P – Eigen vector centrality for PTSD subjects, Eig_C – Eigen vector centrality for control subjects, p_degree – p value for degree centrality comparison, p_betweenness – p value for betweenness centrality comparison, p_closeness – p value for closeness centrality comparison, p_eigenvector – p value for eigenvector centrality comparison.

Supplemental Table 11- Cortical Thickness - Positive Correlations

50-60 years

| PTSD > CONT | | | | | | | | |
| --- | --- | --- | --- | --- | --- | --- | --- | --- |
| Nodes | Deg_P | Deg_C | Bet_P | Bet_C | Clo_P | Clo_C | Eig_P | Eig_C |
| L_S_postcentral | 31.00 | 22.00 | 664.51 | 154.21 | 81.33 | 74.67 | 0.21 | 0.17 |
| R_G_insular_short | 15.00 | 6.00 | 355.94 | 80.01 | 70.75 | 55.25 | 0.02 | 0.002 |
| CONT > PTSD | | | | | | | | |
| R_S_oc_temp_med_and_Lingual | 7.00 | 21.00 | 64.99 | 1064.00 | 59.08 | 76.83 | 0.01 | 0.05 |
| R_S_orbital_lateral | 7.00 | 18.00 | 1.98 | 252.39 | 58.58 | 69.17 | 0.03 | 0.03 |
| R_S_orbital_H_Shaped | 8.00 | 17.00 | 37.66 | 352.39 | 61.25 | 70.92 | 0.01 | 0.03 |
| L_G_precuneus | 17.00 | 30.00 | 96.38 | 390.12 | 71.25 | 81.17 | 0.12 | 0.22 |
| L_S_parieto_occipital | 17.00 | 29.00 | 127.58 | 467.37 | 70.75 | 79.83 | 0.11 | 0.20 |

| PTSD > CONT | | | |
| --- | --- | --- | --- |
| p_degree | p_betweenness | p_closeness | p_eigenvector |
| 0.05 | 0.003 | 0.05 | 0.09 |
| 0.02 | 0.04 | 0.01 | 0.03 |
| CONT > PTSD | | | |
| 0.0008 | 0.0008 | 0.0006 | 0.01 |
| 0.02 | 0.02 | 0.01 | 0.45 |
| 0.04 | 0.04 | 0.01 | 0.05 |
| 0.03 | 0.02 | 0.01 | 0.02 |
| 0.02 | 0.01 | 0.01 | 0.03 |

Deg_P – Degree centrality for PTSD subjects, Deg_C – Degree centrality for control subjects, Bet_P – Betweenness centrality for PTSD subjects, Bet_C – Betweenness centrality for control subjects, Clo_P – Closeness centrality for PTSD subjects, Clo_C – Closeness centrality for control subjects, Eig_P – Eigen vector centrality for PTSD subjects, Eig_C – Eigen vector centrality for control subjects, p_degree – p value for degree centrality comparison, p_betweenness – p value for betweenness centrality comparison, p_closeness – p value for closeness centrality comparison, p_eigenvector – p value for eigenvector centrality comparison.

Supplemental Table 12- Cortical Thickness - Positive Correlations

>60 years

| PTSD > CONT | | | | | | | | |
| --- | --- | --- | --- | --- | --- | --- | --- | --- |
| Nodes | Deg_P | Deg_C | Bet_P | Bet_C | Clo_P | Clo_C | Eig_P | Eig_C |
| L_G_temporal_inf | 27.00 | 9.00 | 571.10 | 64.86 | 74.75 | 57.83 | 0.04 | 0.01 |
| L_S_oc_temp_med_and_Lingual | 25.00 | 13.00 | 512.83 | 138.41 | 72.50 | 62.25 | 0.03 | 0.01 |
| L_S_precentral_inf_part | 26.00 | 9.00 | 925.08 | 182.36 | 79.58 | 64.45 | 0.14 | 0.04 |
| CONT > PTSD | | | | | | | | |
| R_S_cingul_marginalis | 9.00 | 25.00 | 17.14 | 186.69 | 63.75 | 74.67 | 0.07 | 0.18 |
| R_S_interm_prim_Jensen | 3.00 | 16.00 | 4.80 | 327.12 | 55.25 | 70.17 | 0.02 | 0.08 |
| R_S_postcentral | 16.00 | 31.00 | 47.50 | 776.19 | 67.50 | 79.75 | 0.13 | 0.20 |
| R_S_subparietal | 4.00 | 16.00 | 91.45 | 29.51 | 56.75 | 67.42 | 0.01 | 0.12 |
| L_G_orbital | 9.00 | 17.00 | 149.42 | 934.16 | 61.08 | 70.67 | 0.01 | 0.01 |
| L_S_oc_sup_and_transversal | 12.00 | 30.00 | 99.46 | 289.76 | 65.25 | 77.33 | 0.06 | 0.20 |

| PTSD > CONT | | | |
| --- | --- | --- | --- |
| p_degree | p_betweenness | p_closeness | p_eigenvector |
| 0.0003 | 0.02 | 0.001 | 0.15 |
| 0.01 | 0.03 | 0.01 | 0.21 |
| 0.01 | 0.005 | 0.01 | 0.02 |
| CONT > PTSD | | | |
| 0.02 | 0.22 | 0.01 | 0.02 |
| 0.01 | 0.07 | 0.01 | 0.04 |
| 0.05 | 0.004 | 0.02 | 0.16 |
| 0.01 | 0.38 | 0.03 | 0.002 |
| 0.04 | 0.01 | 0.02 | 0.38 |
| 0.02 | 0.18 | 0.01 | 0.01 |

Deg_P – Degree centrality for PTSD subjects, Deg_C – Degree centrality for control subjects, Bet_P – Betweenness centrality for PTSD subjects, Bet_C – Betweenness centrality for control subjects, Clo_P – Closeness centrality for PTSD subjects, Clo_C – Closeness centrality for control subjects, Eig_P – Eigen vector centrality for PTSD subjects, Eig_C – Eigen vector centrality for control subjects, p_degree – p value for degree centrality comparison, p_betweenness – p value for betweenness centrality comparison, p_closeness – p value for closeness centrality comparison, p_eigenvector – p value for eigenvector centrality comparison.

Supplemental Table 13- Surface Area – Negative Correlations

| PTSD > CONT | | | | | | | | |
| --- | --- | --- | --- | --- | --- | --- | --- | --- |
| Nodes | Deg_P | Deg_C | Bet_P | Bet_C | Clo_P | Clo_C | Eig_P | Eig_C |
| L_G_occipital_middle | 21.00 | 9.00 | 180.58 | 36.60 | 78.50 | 69.17 | 0.09 | 0.04 |
| L_S_collat_transv_ant | 29.00 | 16.00 | 541.71 | 122.89 | 83.83 | 74.67 | 0.09 | 0.06 |
| L_S_front_sup | 37.00 | 27.00 | 631.91 | 245.83 | 90.67 | 85.50 | 0.17 | 0.14 |
| R_G_and_S_cingul_mid_Ant | 16.00 | 7.00 | 122.73 | 23.00 | 77.67 | 67.08 | 0.08 | 0.03 |
| R_G_front_inf_Triangul | 15.00 | 5.00 | 119.03 | 18.01 | 76.33 | 64.33 | 0.06 | 0.02 |
| CONT > PTSD | | | | | | | | |
| R_G_pariet_inf_Supramar | 20.00 | 34.00 | 227.53 | 650.21 | 79.42 | 89.50 | 0.10 | 0.15 |
| R_G_parietal_sup | 16.00 | 30.00 | 128.15 | 544.06 | 74.83 | 85.25 | 0.06 | 0.11 |
| R_G_rectus | 2.00 | 8.00 | 4.55 | 29.24 | 52.33 | 68.08 | 0.003 | 0.04 |
| R_S_circular_insula_inf | 7.00 | 16.00 | 26.58 | 248.57 | 66.08 | 79.50 | 0.03 | 0.07 |
| R_S_front_middle | 14.00 | 23.00 | 191.76 | 623.90 | 73.42 | 79.00 | 0.04 | 0.06 |
| L_G_front_inf_Triangul | 15.00 | 24.00 | 115.68 | 280.56 | 77.17 | 82.17 | 0.07 | 0.10 |
| L_G_temp_sup_Plan_polar | 2.00 | 6.00 | 2.32 | 22.98 | 53.33 | 66.25 | 0.01 | 0.03 |

| PTSD > CONT | | | |
| --- | --- | --- | --- |
| p_degree | p_betweenness | p_closeness | p_eigenvector |
| 0.01 | 0.06 | 0.01 | 0.01 |
| 0.004 | 0.01 | 0.01 | 0.01 |
| 0.005 | 0.001 | 0.01 | 0.02 |
| 0.03 | 0.07 | 0.01 | 0.02 |
| 0.01 | 0.06 | 0.01 | 0.02 |
| CONT > PTSD | | | |
| 0.01 | 0.05 | 0.002 | 0.01 |
| 0.01 | 0.01 | 0.004 | 0.02 |
| 0.04 | 0.04 | 0.03 | 0.02 |
| 0.02 | 0.01 | 0.003 | 0.04 |
| 0.05 | 0.03 | 0.08 | 0.03 |
| 0.02 | 0.04 | 0.06 | 0.05 |
| 0.04 | 0.02 | 0.02 | 0.01 |

Deg_P – Degree centrality for PTSD subjects, Deg_C – Degree centrality for control subjects, Bet_P – Betweenness centrality for PTSD subjects, Bet_C – Betweenness centrality for control subjects, Clo_P – Closeness centrality for PTSD subjects, Clo_C – Closeness centrality for control subjects, Eig_P – Eigen vector centrality for PTSD subjects, Eig_C – Eigenvector centrality for control subjects, p_degree – p value for degree centrality comparison, p_betweenness – p value for betweenness centrality comparison, p_closeness – p value for closeness centrality comparison, p_eigenvector – p value for eigenvector centrality comparison.

Supplemental Table 14- Surface Area - Positive Correlations (<10 years)

| PTSD > CONT | | | | | | | | |
| --- | --- | --- | --- | --- | --- | --- | --- | --- |
| Nodes | Deg_P | Deg_C | Bet_P | Bet_C | Clo_P | Clo_C | Eig_P | Eig_C |
| L_G_insular_short | 25.00 | 15.00 | 581.34 | 196.50 | 76.33 | 69.83 | 0.22 | 0.14 |
| L_G_temp_sup_Plan_tempo | 23.00 | 11.00 | 563.49 | 242.82 | 75.50 | 66.33 | 0.20 | 0.06 |
| L_Pole_temporal | 19.00 | 9.00 | 536.54 | 134.57 | 72.67 | 63.25 | 0.15 | 0.03 |
| L_S_circular_insula_ant | 26.00 | 13.00 | 554.78 | 224.50 | 77.25 | 69.00 | 0.22 | 0.10 |
| CONT > PTSD | | | | | | | | |
| L_S_intrapariet_and_P_trans | 3.00 | 11.00 | 14.97 | 150.87 | 47.65 | 64.08 | 0.001 | 0.04 |
| L_S_parieto_occipital | 5.00 | 17.00 | 10.57 | 362.83 | 53.95 | 73.33 | 0.01 | 0.14 |
| R_G_and_S_subcentral | 3.00 | 14.00 | 15.53 | 431.09 | 49.03 | 72.33 | 0.002 | 0.09 |
| R_G_and_S_cingul_mid_Post | 5.00 | 13.00 | 55.95 | 148.04 | 55.17 | 66.58 | 0.004 | 0.11 |
| R_G_pariet_inf_Angular | 6.00 | 14.00 | 165.10 | 309.36 | 56.25 | 67.17 | 0.01 | 0.07 |
| R_G_temp_sup_Plan_polar | 3.00 | 10.00 | 23.45 | 247.55 | 49.95 | 66.67 | 0.004 | 0.05 |
| R_S_central | 4.00 | 12.00 | 37.16 | 303.65 | 54.50 | 68.25 | 0.005 | 0.06 |
| R_S_interm_prim_Jensen | 6.00 | 11.00 | 196.90 | 214.94 | 58.75 | 66.08 | 0.01 | 0.06 |
| R_S_suborbital | 4.00 | 14.00 | 27.21 | 234.50 | 51.78 | 69.92 | 0.01 | 0.13 |
| L_G_subcallosal | 10.00 | 20.00 | 133.01 | 526.40 | 61.00 | 76.00 | 0.01 | 0.17 |
| L_G_temporal_middle | 5.00 | 15.00 | 51.15 | 293.42 | 55.03 | 71.67 | 0.01 | 0.12 |
| L_Lat_fis_ant_Vertical | 6.00 | 15.00 | 57.06 | 453.91 | 59.25 | 74.17 | 0.02 | 0.12 |

| PTSD > CONT | | | |
| --- | --- | --- | --- |
| p_degree | p_betweenness | p_closeness | p_eigenvector |
| 0.02 | 0.03 | 0.04 | 0.04 |
| 0.02 | 0.06 | 0.04 | 0.01 |
| 0.04 | 0.02 | 0.08 | 0.03 |
| 0.004 | 0.04 | 0.03 | 0.02 |
| CONT > PTSD | | | |
| 0.03 | 0.11 | 0.003 | 0.04 |
| 0.04 | 0.01 | 0.004 | 0.09 |
| 0.002 | 0.002 | 0.0004 | 0.002 |
| 0.03 | 0.23 | 0.02 | 0.003 |
| 0.03 | 0.18 | 0.04 | 0.008 |
| 0.01 | 0.02 | 0.004 | 0.02 |
| 0.01 | 0.02 | 0.01 | 0.01 |
| 0.02 | 0.19 | 0.05 | 0.02 |
| 0.01 | 0.06 | 0.003 | 0.004 |
| 0.06 | 0.01 | 0.03 | 0.01 |
| 0.02 | 0.05 | 0.005 | 0.004 |
| 0.03 | 0.005 | 0.01 | 0.04 |

Deg_P – Degree centrality for PTSD subjects, Deg_C – Degree centrality for control subjects, Bet_P – Betweenness centrality for PTSD subjects, Bet_C – Betweenness centrality for control subjects, Clo_P – Closeness centrality for PTSD subjects, Clo_C – Closeness centrality for control subjects, Eig_P – Eigenvector centrality for PTSD subjects, Eig_C – Eigen vector centrality for control subjects, p_degree – p value for degree centrality comparison, p_betweenness – p value for betweenness centrality comparison, p_closeness – p value for closeness centrality comparison, p_eigenvector – p value for eigenvector centrality comparison.

Supplementary Table 15- Surface Area- Positive Correlations -10-15 years

| PTSD > CONT | | | | | | | | |
| --- | --- | --- | --- | --- | --- | --- | --- | --- |
| Nodes | Deg_P | Deg_C | Bet_P | Bet_C | Clo_P | Clo_C | Eig_P | Eig_C |
| R_S_oc_middle_and_Lunatus | 18.00 | 7.00 | 427.18 | 38.23 | 72.58 | 59.33 | 0.13 | 0.03 |
| R_Lat_fis_post | 21.00 | 12.00 | 514.23 | 120.88 | 75.92 | 67.33 | 0.18 | 0.11 |
| CONT > PTSD | | | | | | | | |
| L_G_and_S_subcentral | 4.00 | 10.00 | 19.85 | 199.40 | 54.08 | 66.08 | 0.02 | 0.03 |
| L_G_and_S_cingul_mid_Post | 4.00 | 13.00 | 24.79 | 320.38 | 56.92 | 70.83 | 0.03 | 0.05 |
| L_G_cingul_Post_ventral | 2.00 | 8.00 | 15.28 | 210.90 | 49.42 | 63.25 | 0.01 | 0.03 |
| L_G_front_inf_Triangul | 10.00 | 26.00 | 122.98 | 539.28 | 64.58 | 77.92 | 0.05 | 0.26 |
| R_S_central | 4.00 | 8.00 | 12.43 | 105.16 | 56.33 | 62.17 | 0.03 | 0.03 |

| PTSD > CONT | | | |
| --- | --- | --- | --- |
| p_degree | p_betweenness | p_closeness | p_eigenvector |
| 0.01 | 0.01 | 0.004 | 0.02 |
| 0.02 | 0.02 | 0.02 | 0.05 |
| CONT > PTSD | | | |
| 0.02 | 0.03 | 0.01 | 0.07 |
| 0.01 | 0.02 | 0.004 | 0.16 |
| 0.01 | 0.03 | 0.003 | 0.07 |
| 0.03 | 0.04 | 0.02 | 0.01 |
| 0.03 | 0.05 | 0.04 | 0.19 |

Deg_P – Degree centrality for PTSD subjects, Deg_C – Degree centrality for control subjects, Bet_P – Betweenness centrality for PTSD subjects, Bet_C – Betweenness centrality for control subjects, Clo_P – Closeness centrality for PTSD subjects, Clo_C – Closeness centrality for control subjects, Eig_P – Eigen vector centrality for PTSD subjects, Eig_C – Eigen vector centrality for control subjects, p_degree – p value for degree centrality comparison, p_betweenness – p value for betweenness centrality comparison, p_closeness – p value for closeness centrality comparison, p_eigenvector – p value for eigenvector centrality comparison.

Supplemental Table 16- Surface Area - Positive Correlations -15-20 years

| PTSD > CONT | | | | | | | | |
| --- | --- | --- | --- | --- | --- | --- | --- | --- |
| Nodes | Deg_P | Deg_C | Bet_P | Bet_C | Clo_P | Clo_C | Eig_P | Eig_C |
| R_Pole_occipital | 19.00 | 10.00 | 317.27 | 101.34 | 71.92 | 60.08 | 0.20 | 0.05 |
| L_G_temp_sup_Lateral | 19.00 | 9.00 | 592.51 | 94.96 | 75.00 | 62.42 | 0.12 | 0.06 |
| L_Pole_occipital | 17.00 | 7.00 | 497.30 | 89.40 | 72.58 | 59.75 | 0.16 | 0.03 |
| L_S_circular_insula_sup | 21.00 | 13.00 | 574.69 | 180.47 | 75.83 | 66.67 | 0.15 | 0.11 |
| L_G_cuneus | 21.00 | 14.00 | 345.51 | 198.45 | 72.92 | 65.17 | 0.21 | 0.07 |
| CONT > PTSD | | | | | | | | |
| L_G_insular_short | 7.00 | 17.00 | 26.27 | 665.57 | 59.50 | 74.00 | 0.05 | 0.11 |
| L_G_precentral | 9.00 | 20.00 | 127.25 | 532.73 | 63.00 | 74.83 | 0.05 | 0.16 |
| L_G_rectus | 3.00 | 19.00 | 7.00 | 780.99 | 56.75 | 75.67 | 0.02 | 0.14 |
| L_S_precentral_sup_part | 9.00 | 22.00 | 88.43 | 495.68 | 61.50 | 74.75 | 0.04 | 0.20 |

| PTSD > CONT | | | |
| --- | --- | --- | --- |
| p_degree | p_betweenness | p_closeness | p_eigenvector |
| 0.02 | 0.16 | 0.02 | 0.002 |
| 0.01 | 0.003 | 0.002 | 0.06 |
| 0.02 | 0.02 | 0.02 | 0.005 |
| 0.03 | 0.01 | 0.01 | 0.13 |
| 0.01 | 0.23 | 0.04 | 0.01 |
| CONT > PTSD | | | |
| 0.03 | 0.001 | 0.01 | 0.10 |
| 0.005 | 0.01 | 0.01 | 0.001 |
| 0.0002 | 0.0002 | 0.0002 | 0.002 |
| 0.003 | 0.01 | 0.01 | 0.0004 |

Deg_P – Degree centrality for PTSD subjects, Deg_C – Degree centrality for control subjects, Bet_P – Betweenness centrality for PTSD subjects, Bet_C – Betweenness centrality for control subjects, Clo_P – Closeness centrality for PTSD subjects, Clo_C – Closeness centrality for control subjects, Eig_P – Eigen vector centrality for PTSD subjects, Eig_C – Eigen vector centrality for control subjects, p_degree – p value for degree centrality comparison, p_betweenness – p value for betweenness centrality comparison, p_closeness – p value for closeness centrality comparison, p_eigenvector – p value for eigenvector centrality comparison.

Supplemental Table 17- Surface Area - Positive Correlations -20-30 years

| PTSD > CONT | | | | | | | | |
| --- | --- | --- | --- | --- | --- | --- | --- | --- |
| Nodes | Deg_P | Deg_C | Bet_P | Bet_C | Clo_P | Clo_C | Eig_P | Eig_C |
| L_S_collat_transv_ant | 10.00 | 4.00 | 339.91 | 22.83 | 60.25 | 48.98 | 0.02 | 0.01 |
| R_G_temp_sup_Plan_polar | 10.00 | 2.00 | 427.11 | 3.47 | 64.25 | 49.93 | 0.05 | 0.01 |
| R_Pole_occipital | 20.00 | 13.00 | 468.40 | 101.85 | 69.17 | 59.00 | 0.11 | 0.01 |
| R_S_cingul_marginalis | 10.00 | 3.00 | 211.67 | 0 | 65.50 | 44.27 | 0.07 | 0.0004 |
| R_S_front_middle | 13.00 | 5.00 | 422.34 | 39.05 | 63.95 | 45.97 | 0.03 | 0.0004 |
| CONT > PTSD | | | | | | | | |
| L_G_Ins_lg_and_S_cent_ins | 8.00 | 24.00 | 60.44 | 549.94 | 60.75 | 73.42 | 0.07 | 0.20 |
| L_S_circular_insula_ant | 16.00 | 32.00 | 221.62 | 467.23 | 68.50 | 78.58 | 0.17 | 0.25 |

| PTSD > CONT | | | |
| --- | --- | --- | --- |
| p_degree | p_betweenness | p_closeness | p_eigenvector |
| 0.01 | 0.04 | 0.04 | 0.10 |
| 0.04 | 0.02 | 0.03 | 0.18 |
| 0.02 | 0.02 | 0.03 | 0.04 |
| 0.05 | 0.08 | 0.03 | 0.02 |
| 0.01 | 0.04 | 0.02 | 0.01 |
| CONT > PTSD | | | |
| 0.01 | 0.04 | 0.02 | 0.02 |
| 0.01 | 0.14 | 0.01 | 0.03 |

Deg_P – Degree centrality for PTSD subjects, Deg_C – Degree centrality for control subjects, Bet_P – Betweenness centrality for PTSD subjects, Bet_C – Betweenness centrality for control subjects, Clo_P – Closeness centrality for PTSD subjects, Clo_C – Closeness centrality for control subjects, Eig_P – Eigen vector centrality for PTSD subjects, Eig_C – Eigen vector centrality for control subjects, p_degree – p value for degree centrality comparison, p_betweenness – p value for betweenness centrality comparison, p_closeness – p value for closeness centrality comparison, p_eigenvector – p value for eigenvector centrality comparison.

Supplemental Table 18- Surface Area - Positive Correlations -30-40 years

| PTSD > CONT | | | | | | | | |
| --- | --- | --- | --- | --- | --- | --- | --- | --- |
| Nodes | Deg_P | Deg_C | Bet_P | Bet_C | Clo_P | Clo_C | Eig_P | Eig_C |
| R_S_temporal_inf | 11.00 | 6.00 | 320.13 | 42.91 | 64.33 | 50.88 | 0.02 | 0.003 |
| L_G_occipital_middle | 14.00 | 5.00 | 570.11 | 55.21 | 65.17 | 49.95 | 0.02 | 0.002 |
| L_Lat_fis_ant_Vertical | 22.00 | 11.00 | 835.28 | 104.72 | 74.08 | 61.53 | 0.19 | 0.12 |
| L_Pole_occipital | 18.00 | 12.00 | 1050.80 | 55.23 | 71.75 | 56.17 | 0.05 | 0.01 |
| L_S_orbital_lateral | 13.00 | 3.00 | 699.13 | 4.92 | 66.33 | 49.98 | 0.08 | 0.01 |
| R_G_and_S_paracentral | 20.00 | 9.00 | 709.87 | 384.04 | 72.92 | 60.50 | 0.08 | 0.01 |
| R_G_and_S_cingul_mid_Post | 21.00 | 12.00 | 689.12 | 195.95 | 74.08 | 63.08 | 0.10 | 0.02 |
| R_G_occipital_middle | 10.00 | 4.00 | 427.85 | 48.88 | 61.42 | 47.28 | 0.02 | 0.001 |
| CONT > PTSD | | | | | | | | |
| L_G_occipital_sup | 6.00 | 14.00 | 107.25 | 538.59 | 54.08 | 61.00 | 0.01 | 0.01 |
| L_G_rectus | 4.00 | 12.00 | 50.89 | 441.86 | 51.45 | 66.75 | 0.01 | 0.04 |
| L_Lat_fis_ant_Horizont | 9.00 | 18.00 | 47.97 | 991.76 | 61.25 | 71.67 | 0.09 | 0.15 |
| L_S_circular_insula_sup | 24.00 | 36.00 | 672.71 | 1208.50 | 73.75 | 81.42 | 0.22 | 0.31 |

| PTSD > CONT | | | |
| --- | --- | --- | --- |
| p_degree | p_betweenness | p_closeness | p_eigenvector |
| 0.02 | 0.02 | 0.001 | 0.04 |
| 0.0002 | 0.002 | 0.001 | 0.02 |
| 0.02 | 0.03 | 0.01 | 0.08 |
| 0.04 | 0.004 | 0.002 | 0.09 |
| 0.02 | 0.003 | 0.01 | 0.06 |
| 0.002 | 0.08 | 0.01 | 0.001 |
| 0.01 | 0.06 | 0.01 | 0.02 |
| 0.01 | 0.01 | 0.004 | 0.03 |
| CONT > PTSD | | | |
| 0.01 | 0.02 | 0.04 | 0.44 |
| 0.03 | 0.03 | 0.01 | 0.10 |
| 0.03 | 0.0002 | 0.02 | 0.12 |
| 0.02 | 0.13 | 0.03 | 0.01 |

Deg_P – Degree centrality for PTSD subjects, Deg_C – Degree centrality for control subjects, Bet_P – Betweenness centrality for PTSD subjects, Bet_C – Betweenness centrality for control subjects, Clo_P – Closeness centrality for PTSD subjects, Clo_C – Closeness centrality for control subjects, Eig_P – Eigen vector centrality for PTSD subjects, Eig_C – Eigen vector centrality for control subjects, p_degree – p value for degree centrality comparison, p_betweenness – p value for betweenness centrality comparison, p_closeness – p value for closeness centrality comparison, p_eigenvector – p value for eigenvector centrality comparison.

Supplemental Table 19- Surface Area - Positive Correlations - 40-50 years

| PTSD > CONT | | | | | | | | |
| --- | --- | --- | --- | --- | --- | --- | --- | --- |
| Nodes | Deg_P | Deg_C | Bet_P | Bet_C | Clo_P | Clo_C | Eig_P | Eig_C |
| R_S_central | 13.00 | 4.00 | 357.26 | 23.59 | 68.00 | 49.90 | 0.09 | 0.01 |
| R_G_temporal_middle | 13.00 | 7.00 | 292.08 | 45.03 | 69.25 | 53.07 | 0.07 | 0.004 |
| L_S_oc_temp_med_and_Lingual | 13.00 | 3.00 | 459.92 | 10.83 | 70.33 | 51.08 | 0.05 | 0.004 |
| L_S_precentral_sup_part | 14.00 | 6.00 | 271.50 | 88.85 | 69.08 | 53.58 | 0.10 | 0.01 |
| L_S_temporal_sup | 18.00 | 11.00 | 628.47 | 475.47 | 72.83 | 62.75 | 0.09 | 0.01 |
| CONT > PTSD | | | | | | | | |
| R_G_temp_sup_Lateral | 7.00 | 18.00 | 86.58 | 681.57 | 60.42 | 70.67 | 0.05 | 0.13 |
| R_Lat_fis_ant_Vertical | 7.00 | 21.00 | 173.75 | 802.52 | 61.17 | 73.25 | 0.05 | 0.21 |
| L_Lat_fis_ant_Horizont | 8.00 | 16.00 | 50.13 | 175.69 | 59.75 | 66.83 | 0.08 | 0.18 |
| L_S_orbital_H_Shaped | 9.00 | 19.00 | 37.45 | 860.43 | 63.42 | 72.50 | 0.08 | 0.11 |

| PTSD > CONT | | | |
| --- | --- | --- | --- |
| p_degree | p_betweenness | p_closeness | p_eigenvector |
| 0.01 | 0.07 | 0.003 | 0.01 |
| 0.04 | 0.09 | 0.001 | 0.01 |
| 0.002 | 0.01 | 0.0004 | 0.02 |
| 0.03 | 0.12 | 0.01 | 0.004 |
| 0.04 | 0.20 | 0.02 | 0.01 |
| CONT > PTSD | | | |
| 0.01 | 0.03 | 0.02 | 0.08 |
| 0.001 | 0.01 | 0.01 | 0.004 |
| 0.03 | 0.11 | 0.02 | 0.02 |
| 0.01 | 0.0008 | 0.03 | 0.21 |

Deg_P – Degree centrality for PTSD subjects, Deg_C – Degree centrality for control subjects, Bet_P – Betweenness centrality for PTSD subjects, Bet_C – Betweenness centrality for control subjects, Clo_P – Closeness centrality for PTSD subjects, Clo_C – Closeness centrality for control subjects, Eig_P – Eigen vector centrality for PTSD subjects, Eig_C – Eigen vector centrality for control subjects, p_degree – p value for degree centrality comparison, p_betweenness – p value for betweenness centrality comparison, p_closeness – p value for closeness centrality comparison, p_eigenvector – p value for eigenvector centrality comparison.

Supplemental Table 20- Surface Area - Positive Correlations- 50-60 years

| PTSD > CONT | | | | | | | | |
| --- | --- | --- | --- | --- | --- | --- | --- | --- |
| Nodes | Deg_P | Deg_C | Bet_P | Bet_C | Clo_P | Clo_C | Eig_P | Eig_C |
| L_S_pericallosal | 17.00 | 6.00 | 439.67 | 131.62 | 72.08 | 61.25 | 0.09 | 0.03 |
| R_G_occipital_sup | 16.00 | 7.00 | 302.49 | 20.52 | 66.58 | 56.92 | 0.10 | 0.03 |
| CONT > PTSD | | | | | | | | |
| R_G_pariet_inf_Supramar | 10.00 | 17.00 | 123.03 | 495.00 | 64.25 | 71.25 | 0.07 | 0.15 |
| L_G_oc_temp_lat_fusifor | 8.00 | 14.00 | 78.32 | 537.32 | 59.17 | 70.33 | 0.03 | 0.06 |
| L_Lat_fis_ant_Vertical | 7.00 | 16.00 | 16.61 | 309.41 | 59.25 | 71.33 | 0.08 | 0.15 |

| PTSD > CONT | | | |
| --- | --- | --- | --- |
| p_degree | p_betweenness | p_closeness | p_eigenvector |
| 0.01 | 0.04 | 0.04 | 0.10 |
| 0.01 | 0.05 | 0.02 | 0.21 |
| CONT > PTSD | | | |
| 0.04 | 0.03 | 0.04 | 0.03 |
| 0.03 | 0.004 | 0.01 | 0.08 |
| 0.04 | 0.02 | 0.003 | 0.25 |

Deg_P – Degree centrality for PTSD subjects, Deg_C – Degree centrality for control subjects, Bet_P – Betweenness centrality for PTSD subjects, Bet_C – Betweenness centrality for control subjects, Clo_P – Closeness centrality for PTSD subjects, Clo_C – Closeness centrality for control subjects, Eig_P – Eigen vector centrality for PTSD subjects, Eig_C – Eigen vector centrality for control subjects, p_degree – p value for degree centrality comparison, p_betweenness – p value for betweenness centrality comparison, p_closeness – p value for closeness centrality comparison, p_eigenvector – p value for eigenvector centrality comparison.

Supplemental Table 21- Surface Area - Positive Correlations -greater than 60 years

| PTSD > CONT | | | | | | | | |
| --- | --- | --- | --- | --- | --- | --- | --- | --- |
| Nodes | Deg_P | Deg_C | Bet_P | Bet_C | Clo_P | Clo_C | Eig_P | Eig_C |
| R_S_temporal_transverse | 17.00 | 7.00 | 172.38 | 107.66 | 70.92 | 61.08 | 0.18 | 0.04 |
| L_G_temporal_middle | 18.00 | 8.00 | 653.26 | 157.24 | 72.75 | 61.42 | 0.06 | 0.04 |
| R_G_Ins_lg_and_S_cent_ins | 22.00 | 8.00 | 514.16 | 74.40 | 75.75 | 61.58 | 0.21 | 0.07 |
| R_G_pariet_inf_Supramar | 22.00 | 12.00 | 661.11 | 435.63 | 76.92 | 69.67 | 0.20 | 0.07 |
| R_G_temp_sup_G_T_transv | 21.00 | 13.00 | 308.56 | 174.34 | 73.83 | 66.58 | 0.22 | 0.10 |
| R_G_temp_sup_Plan_tempo | 21.00 | 6.00 | 321.69 | 83.17 | 74.58 | 57.92 | 0.22 | 0.03 |
| R_Lat_fis_post | 18.00 | 9.00 | 332.06 | 125.46 | 71.92 | 64.75 | 0.19 | 0.08 |
| CONT > PTSD | | | | | | | | |
| R_S_cingul_marginalis | 9.00 | 15.00 | 120.07 | 403.90 | 63.33 | 71.67 | 0.03 | 0.11 |
| R_S_collat_transv_post | 2.00 | 13.00 | 6.61 | 187.85 | 49.33 | 66.75 | 0.004 | 0.09 |
| R_S_precentral_sup_part | 4.00 | 13.00 | 51.58 | 158.21 | 56.25 | 67.25 | 0.01 | 0.10 |
| R_G_oc_temp_med_Lingual | 11.00 | 20.00 | 219.08 | 647.38 | 63.00 | 73.08 | 0.03 | 0.13 |

| PTSD > CONT | | | |
| --- | --- | --- | --- |
| p_degree | p_betweenness | p_closeness | p_eigenvector |
| 0.02 | 0.36 | 0.03 | 0.01 |
| 0.01 | 0.005 | 0.02 | 0.26 |
| 0.002 | 0.02 | 0.004 | 0.01 |
| 0.02 | 0.11 | 0.05 | 0.01 |
| 0.02 | 0.22 | 0.05 | 0.00 |
| 0.003 | 0.11 | 0.001 | 0.001 |
| 0.02 | 0.16 | 0.04 | 0.02 |
| CONT > PTSD | | | |
| 0.05 | 0.03 | 0.05 | 0.01 |
| 0.03 | 0.13 | 0.003 | 0.01 |
| 0.03 | 0.14 | 0.04 | 0.01 |
| 0.04 | 0.04 | 0.03 | 0.01 |

Deg_P – Degree centrality for PTSD subjects, Deg_C – Degree centrality for control subjects, Bet_P – Betweenness centrality for PTSD subjects, Bet_C – Betweenness centrality for control subjects, Clo_P – Closeness centrality for PTSD subjects, Clo_C – Closeness centrality for control subjects, Eig_P – Eigen vector centrality for PTSD subjects, Eig_C – Eigen vector centrality for control subjects, p_degree – p value for degree centrality comparison, p_betweenness – p value for betweenness centrality comparison, p_closeness – p value for closeness centrality comparison, p_eigenvector – p value for eigenvector centrality comparison.

Supplementary table 22- Compilation of overlapping regions of interest - CT

| Brain Areas | CT Positive Corr | Females | <10 years | 10-20 years | 20-30 years | Sun et al 2018 (Children) | Sun et al 2018 (Youth) | Sun et al 2018 (Veterans) | Mueller et al 2015 |
| --- | --- | --- | --- | --- | --- | --- | --- | --- | --- |
| L-precentral | P > C | P>C (M) | C > P |  |  |  |  | R<C, A>R, A>C | |
| L-anterior cingulate |  |  |  | P >C |  | P>C | P > Ma, P>C | |  |
| L-posterior cingulate |  | F>M (P) |  |  |  |  |  | R>C, C>A |  |
| L-fusiform | P > C |  |  |  |  |  |  |  |  |
| L-inferior frontal | P > C |  |  |  |  | Ma>P, C>Ma, C>P |  |  |  |
| L-paracentral |  |  | C>P |  |  |  |  |  |  |
| L-insula |  | F>M(P), C>P(M),P>C(F) |  |  |  |  |  |  | Combined thickness (left plus right) decreased in PTSD positive compared to PTSD negative |
| R-insula |  |  |  |  |  | P>C | P>C | R>C |  |
| R-anterior cingulate |  |  |  |  |  |  |  | R>A |  |
| R-posterior cingulate |  |  |  | C>P | P>C | P>C |  | R>A |  |
| R-paracentral | P>C | P>C (M) |  |  | P>C |  |  |  |  |
| R-ant occipital | C >P |  |  | C>P |  |  |  | R>C |  |
| R-rectus |  | M>F (P), P>C(M) |  |  |  |  |  |  |  |
| R-inferior frontal |  |  |  |  |  |  |  | R>C, R>A |  |

P – PTSD subjects, C- control subjects, R – subjects with PTSD in remission, A – subjects with active PTSD, Ma – Maltreated subjects with no PTSD, M- males, F- females.

Supplementary Table 23- Compilation of overlapping regions of interest - SA

| Brain Areas | SA Positive Corr | Females | <10 years | 10-20 years | 20-30 years |
| --- | --- | --- | --- | --- | --- |
| L-precentral |  |  |  | C>P |  |
| L-anterior cingulate |  | M>F (P), P>C (M) |  |  |  |
| L-posterior cingulate |  |  |  | C>P |  |
| L-rectus |  |  |  | C>P |  |
| L-insula |  |  | P>C | C>P |  |
| L-subcallosal |  |  | C>P |  |  |
| R-insula |  | F>M (C) |  |  |  |
| R-anterior cingulate |  |  |  |  |  |
| R-posterior cingulate | P>C |  | C>P |  | P>C |
| R-rectus |  | M>F(C), C>P(M) |  |  |  |
| Brain Areas | SA Positive Corr | Females | <10 years | 10-20 years | 20-30 years |

P – PTSD subjects, C- control subjects, M- males, F- females

Supplementary Table 24– Inclusion and exclusion criteria

| Site | Inclusion criteria | Exclusion criteria |
| --- | --- | --- |
| AMC Amsterdam | All: 18-65 years of age, police officers, eligible for MRI. PTSD: current PTSD diagnosis, with CAPS ≥ 45. Controls: exposure to at least one traumatic event (according to DSM-IV A1 criterion), with CAPS < 15 | All: history of neurological disorders, any severe or chronic systemic disease or unstable medical condition (including endocrinological disorders), use of psychotropic medications. Females: pregnancy or breastfeeding. PTSD: current psychotic disorder, substance-related disorder, severe personality disorder, severe major depressive disorder (MDD) (i.e., involving high suicidal risk and/or psychotic symptoms) or current suicidal risk. Controls: any current Axis-1 disorder and lifetime history of PTSD or MDD |
| Cape Town | Between 18 and 65 years, speak English, Afrikaans or Xhosa | Mental retardation, critical medical condition, current psychotic episode/disorder, contraindications for MRI (e.g., metal objects in body, pacemakers) |
| DoD ADNI | *PTSD*: Subjects must be Veterans of the Vietnam War, 50-90 years of age. Subjects who meet the SCID-I (for DSM-IV-TR) criteria for current/chronic PTSD (identified by records and verified by our telephone assessments). In addition to meeting DSM-IV-TR criteria for current/chronic PTSD, subjects must have a minimum current CAPS score of 50 as determined by telephone assessment. The PTSD symptoms contributing to the PTSD Diagnosis and Current CAPS score must be related to a Vietnam War related trauma. Must live within 150 miles of the closest ADNI clinic in subject’s area. *Control:* Subjects must be Veterans of the Vietnam War, 50-90 years of age. Comparable in age, sex, and education with TBI and PTSD groups May be receiving VA disability payments for something other than TBI or PTSD – or no disability at all. Must live within 150 miles of the closest ADNI clinic in subject’s area | *PTSD*: Mild Cognitive Impairment/Dementia Documented or self-report history of mild/moderate severe TBI Any history of head trauma associated with injury onset cognitive complaints, or Loss of consciousness for >5minutes. Control: MCI/Dementia Presence of PTSD by SCID-I for DSM-IV-TR criteria, or a CAPS score of >30 (Both current and/or a history of PTSD will be excluded). Documented or self-report history of mild/moderate severe TBI Any history of head trauma associated with injury onset cognitive complaints, or Loss of Consciousness for >5 minutes History of PTSD or current PTSD Exclusionary criteria applied to TBI/PTSD will be applied to controls. ALL: MCI/dementia History of psychosis or bipolar affective disorder; History of alcohol or substance abuse/dependence within the past 5 years (by DSM IV – TR criteria); MRI-related exclusions: aneurysm clips, metal implants that are determined to be unsafe for MRI; and/or claustrophobia; Contraindications for lumbar puncture, PET scan, or other procedures in this study; Any major medical condition must be stable for at least 4 months prior to enrollment. These include but are not limited to clinically significant hepatic, renal, pulmonary, metabolic or endocrine disease, cancer, HIV infection and AIDS, as well as cardiovascular disease. Seizure disorder or any systemic illness affecting brain function during the past 5 years will be exclusionary Clinical evidence of stroke. Have a history of relevant severe drug allergy or hypersensitivity. Subjects with current clinically significant unstable medical comorbidities, as indicated by history or physical exam, that pose a potential safety risk to the subject. |
| Duke/Durham VA | 18-65, OEF/OIF veterans, fluent in English, free of implanted metal objects or metal shards in eyes, antidepressant, sleep, and anti-anxiety medication permitted | Axis I other than PTSD or MDD, current substance abuse or lifetime substance dependence (other than nicotine), high risk for suicide, claustrophobia, neurological disorders, learning disability or developmental delay, major medical conditions |
| Emory GTP | 18-65 years of age, endorsed at least 1 criterion A trauma, English-speaking | Current psychotic symptoms or bipolar disorder, current substance or alcohol dependence, history of head trauma, taking any psychoactive medication, current illegal drug use (verified with urine drug screen within 24 hours of scan) |
| McLean | History of childhood maltreatment; Legal and mental competency of the patient; Female; All ethnic backgrounds; Age between 18 and 60; Fluent English speakers; Normal or Corrected Vision | Male; Under 18 or over 60; Delirium secondary to medical illness; History of neurological conditions that may cause significant psychiatric symptomatology (e.g., dementia); Any contraindication to MR scans, including claustrophobia, pregnancy, metal implants, etc.; Current alcohol or substance use disorder (within the last month); A history of schizophrenia or other psychotic disorder; History of head injury or loss of consciousness for longer than 5 min (including concussion); Positive pregnancy test |
| UNSW | 18-60 years | Neurological disorders, under 18; over 60; traumatic brain injury; psychosis |
| U of Sydney | 18-65 years of age, endorsed at least 1 criterion A trauma, English-speaking | History of neurological illness (Huntington’s, Parkinson’s, dementia, MS, etc.). History of Seizure Disorders, unrelated to head injury(is). Current diagnosis of schizophrenia spectrum or other psychotic disorders (not related to PTSD). Current diagnosis of bipolar or related disorders (not related to PTSD). Current active homicidal and/or suicidal ideation with intent requiring crisis intervention. Cognitive disorder due to general medical condition other than TBI. Unstable psychological diagnosis that would interfere with accurate data collection, determined by consensus of at least two doctorate-level psychologists. Also, MRI contra-indications for MRI including metallic implants or foreign objects deemed unsafe by the MRI technician (such as but not limited to pacemaker, shrapnel, metallic screws). Surgery in the past 2 months except as approved by the MRI technician (such as but not limited to dental work, colonoscopy). Weight exceeding the capacity of the scanner table. |
| U of Washington | Aged 8-20 | Psychiatric medication use (excepting stimulant meds for ADHD), braces, claustrophobia, active substance dependence, pervasive developmental disorder, non-English speaking, active safety concerns. |
| VUMC Amsterdam | 18-65 | Antisoc. pers. disorder, DID, recurrent psychoses, current drug abuse or dependence, medication other than stable SSRIs or infrequent benzodiazepine use |
| West Haven VA | combat-exposed Veterans with PTSD and 21 age-matched male combat-exposed healthy controls (combat controls; CC). All participants had been deployed on one or more tours to Iraq and/or Afghanistan and reported exposure to combat-related experiences. All participants were 18 to 50 years of age. | Participants were excluded based on moderate and severe TBI, neurological disorder, and MRI contraindications. Participants with PTSD were also excluded on the basis of a diagnosis of current drug/alcohol abuse, recent change in antidepressant medications (stable dose for 4-weeks required). |
| Yale | Participants ranged in age from 21 to 60 and had been deployed on one or more combat tours. | Individuals were excluded from the study if they met any of the following criteria: a diagnosis of bipolar disorder or psychotic disorder, as assessed by the SCID-IV (First, Spitzer, Gibbon, & Williams, 2002); current benzodiazepine use; a history of ADHD, learning disorder, moderate or severe traumatic brain injury (TBI), brain tumor, epilepsy, or a neurological disorder; current inpatient status; or an MRI contraindication. |
| South Dakota | Military cohort: OEF/OIF  SAP cohort: Participants were undergraduate students who were identified as an adult child of an alcoholic parent (ACoA), based on the Children of Alcoholics Screening Test (CAST, Jones, 1983). A score of 6 or above on the CAST indicated the participant was more than likely the child of an alcoholic/s and raised by this parent/s. | Participants were excluded for current or previous seizure history, contraindications to MRI, or if they exhibited possible psychotic or other psychological symptoms that would make inclusion in the study potentially hazardous to them. |
| Stanford | Study 1: Psychiatric diagnoses, or absence thereof for controls, were based on DSM-IV criteria using the Clinician-Administered PTSD Scale (CAPS) for PTSD and the Structured Clinical Interview for the Diagnostic and Statistical Manual of Mental Disorders Axis I (SCID I) for other Axis I disorders. Participants were permitted to meet diagnostic criteria for comorbid mood and anxiety disorders secondary to PTSD. Trauma-exposed healthy controls were required to have experienced a criterion A trauma, but not meet lifetime criteria for any Axis 1 psychiatric disorder, including PTSD.  Study 2: Similar inclusion and diagnostic criteria were used as above except that diagnoses were based on DSM-5 criteria rather than DSM-IV. | Study 1 and 2: General exclusion criteria for both groups included the following: a history of psychotic, bipolar or substance dependence (within 3 months for patients and lifetime for controls), a history of a neurological disorder, greater than mild traumatic brain injury (i.e. >30 minutes loss of consciousness or >24-hour post-trauma amnesia), claustrophobia, and regular use of benzodiazepines, opiates, thyroid medications, or other CNS medication. |
| Ghent University | For trauma group: experience(s) of physical, sexual, and/or emotional abuse occurring before 17 years of age as per SLESQ. For comparisons: no experience of childhood trauma and no experience of abuse-related trauma (e.g. emotional abuse, physical/sexual assault, etc.) later in life. For both: MRI compatibility (i.e., no pregnancy or metal implants), fluency in Dutch, normal or corrected-to-normal vision, female, and being 18-60 years of age | History of severe head trauma or severe neurological condition |
| Leiden University | All participants met the following inclusion criteria: aged between 12 and 21, estimated full scale IQ (FIQ)≥80 as measured by Dutch versions of the Wechsler Intelligence Scales for Children (WISC-III) or adults (WAIS), being right-handed, normal or corrected-to-normal vision, sufficient understanding of the Dutch language, no history of neurological impairments and no contraindications for MRI testing (e.g. braces, metal implants or possible pregnancy). | Primary DSM-IV diagnosis of ADHD, pervasive developmental disorders, Tourette’s syndrome, obsessive–compulsive disorder, bipolar disorder, and psychotic disorders, current use of psychotropic medication other than stable use of SSRI’s, or amphetamine medication on the day of scanning, and current substance abuse. |
| University of Wisconsin (Larson) | PTSD criterion A met, age 18-60, GCS>= 13 (mild TBI criteria), Rothbaum 3 or higher or item 2 rated 3 or higher, English speaking (either native or bilingual proficiency), able to schedule within 30 days of brain injury. | Still in high school, re-admitted to hospital for current brain injury, live too far away to travel for study, police hold, incarcerated, intentional self-inflicted injury, known perpetrator, moderate to severe cognitive impairment, loss of consciousness > 30 minutes, pregnant, clear evidence of substance abuse, anti-psychotic or anti-seizure medication, indication of psychotic disorder or manic symptoms, MRI contraindications, history of seizures or other neurological conditions, severe hearing or vision problems |
| Minneapolis VA | Participants were veterans of Operation Enduring Freedom and/or Operation Iraqi Freedom, age 22-62, who had been exposed to combat during their deployment(s). | Participants were excluded from the study if they met criteria for 1) a current substance-induced psychotic disorder or psychotic disorder due to a general medical condition (other than TBI), 2) current DSM-IV substance abuse or dependence other than alcohol, caffeine, or nicotine, 3) a moderate or severe traumatic brain injury from either impact or blast, 4) a neurologic condition other than TBI, 5) a current unstable medical condition that would likely affect brain function (e.g., uncontrolled diabetes), or 6) significant imminent risk of suicidal or homicidal behavior. |
| Muenster | All patients fulfilled the diagnostic criteria for PTSD as primary diagnosis according to the DSM-IV-TR (American Psychiatric Association, 2000), assessed by the German version of the Structured Clinical Interview for DSM-IV (SCID; Wittchen et al., 1997). Given the focus on IPV-PTSD, the experience of a trauma related to IPV (e.g., rape, sexual or physical abuse) at least once was an inclusion criterion for the patient group. All participants had normal or corrected-to-normal vision and were right-handed as determined by the Edinburgh Handedness Inventory (Oldfield, 1971). | No control had a lifetime PTSD. MRI contraindications. |
| WACO VA | Study 1: Age 18-60, US military veteran  Study 2: Age 18-60, veteran with a clinical diagnosis of TBI in the VA medical records  Study 3: Age 18-60, veteran enrolled in a residential PTSD treatment program | Study 1: Serious general medication condition that would risk the subject being able to complete MRI (active seizure disorder, dementia, active back or muscle spasms), MRI safety factors (metal in body, claustrophobia, etc) or quality issues (tremors, teeth braces), pregnant or nursing females  Study 2: Absence of qEEG parameters more than 2 standard deviations from the population mean of healthy age-matched historical controls saved in a commercial normative database (Neuroguide, Largo, FL), positive screen on the MINI International Neuropsychiatric Interview[13]: diagnosis of schizophrenia, schizoaffective disorder, bipolar disorder type I, MIR safety factors or quality problems, severe substance use disorder, high risk of suicide, pregnant females  Study 3: MRI safety factors or quality issues, pregnant or nursing females, current psychosis including Axis I psychotic disorder, bipolar disorder, or schizophrenia, demenia or another severe cognitive disorder, prior exposure to TMS, seizure disorder, positive screen for suicidal intent, plan, or behavior within the past 6 months, a TMS motor threshold of 70% or great of the machine’s maximum output |
| Intrust Consortium | Study 1: Deployment during recent OEF/OIF/Operation New Dawn (OND) conflict and meeting DSM-IV diagnostic criteria for one or more anxiety or depressive disorders.  Study 2: Cognitive complaints and either PTSD and/or TBI  Study 3: Veteran or civilian outpatients with a primary diagnosis of PTSD and Clinician Administered PTSD Scale score greater than 49.  Study 4: Returning from OEF/OIF theatre for medical reasons, being a patient at WRAMC or NNMC, diagnosed as TBI positive (with initial score of 13-15 on the Glasgow Coma Scale) or negative using DoD criteria, aged 18-40, and DEERS eligible.  Study 5:English language literacy to provide content, negative pregnancy test for women, Glasgow Coma Scale score of 15, extension of GCS with 7-point amnesia scale score of 6 or 7, and clinically judged to be at low risk for taking tramadol.  Study 6: Age 18-60, active duty or veteran recently screened positive for TBI, receiving care at one the designated study clinical sites, capable of giving informed consent, Defense Enrollment Eligibility Reporting System (DEERS) eligible (subset of participants), verified TBI positive (subset), for those with TBI, GCS score of 13-15. | Study 1: Significant cognitive impairment, severe psychopathology (psychosis, bipolar disorder, imminent suicidality, untreated substance dependence), recent change in pharmacological intervention, concurrent psychotherapy for the target complaint, anticipated change in circumstances that would prevent study completion (e.g., deployment).  Study 2: Prior adverse reaction to or taking medications that would interfere with study medications, women who were or were planning to become pregnant or lactating, certain medical exclusions, inability to meet English language proficiency, or history of psychotic disorder, bipolar I, alcohol or stimulant use disorder, tic disorder, severe depression or acute suicidality.  Study 3: Unstable medical disease or alanine transferase/aspartate transferase levels significant above normal (or abnormal liver function tests), moderate or severe TBI, pregnant or nursing women, history of psychotic, bipolar, or neurodegenerative disease, history of substance use disorder, imminent suicidality, current use of psychotropic medications or recent use of investigational medications, current engagement in evidence-based treatment for PTSD, current litigation for traumatic event, unwillingness to abstain from grapefruit products during the trial, or lack of English proficiency.  Study 4: Non-medical military separation, inability to provide consent (e.g., due to English language proficiency level), history of penetrating head injury, significant neurological conditions, current treatment for illness that could impact brain functioning, history of major psychiatric conditions including drug or alcohol use disorders, IV medication use for pain.  Study 5: Pregnant or nursing, homeless, active suicidal or homicidal with plans/ intent, history of lifetime opioid dependence or abuse, psychosis, other substance abuse or dependence (except tobacco) in the past 60 days, anorexia nervosa, antisocial personality disorder, or other psychiatric conditions more clinically prominent than PTSD, serious or unstable illness or history of stroke, seizures, or brain tumor, use of non-study medications unless approved by PI (e.g., benzodiazepines), current psychotherapy, reaction to tramadol.  Study 6: Speech/language deficit of sufficient severity to preclude answering interview questions, unable or unwilling to provide informed consent and Health Insurance Portability and Accountability Act (HIPAA)authorization ("unable" includes cases in which the potential subject cannot read and understand English well enough to provide informed consent), second level in-depth TBI evaluation done prior to SAFE TBI interview (subset), penetrating head injury (subset), record of drug or alcohol abuse or dependence in the past six months as documented in medical chart (subset), Structured Clinical Interview for Diagnostic and Statistical Manual of Mental Disorders -IV (SCID) current or lifetime PTSD diagnosis related to life events that occurred prior to most recent deployment (subset), taking intravenous medications for pain; participation will be delayed until such medication has been discontinued (subset). |
| McLean (Rosso) | 1) 20-50 years of age; 2) right-handed; 3) DSM-IV diagnosis consistent with group assignment; 4) ability to provide written informed consent. | 1) Medical condition that would confound results; 2) history of seizures or head trauma with loss of consciousness; 3) exposure to psychotropic medications within 4 weeks of study (8 weeks for fluoxetine); 4) metal implants, claustrophobia or other MR exclusions; 5) positive urine toxicology or HCG status on scan day; 6) history of psychotic disorder, bipolar disorder, eating disorder, mental retardation, or pervasive developmental disorder; history of meeting full DSM-IV criteria for non-PTSD anxiety disorder. |
| UCAS | One member in each household was randomly selected as participants; All participants were between 16 and 65 years old during the survey, and all personally experienced the 2008 earthquake | Those with major psychosis (e.g. schizophrenia and organic mental disorders) were excluded. |
| Duke DeBellis | Maltreated: Positive forensic investigation conducted by CPS that indicated physical and sexual abuse and/or neglect. Healthy volunteers: no history of DSM Axis I disorders or Type A traumas, negative maltreatment screen on initial telephone interview, negative history of participant or participant sibling maltreatment or negative review of pediatric and birth medical records that met or would have met state CPS maltreatment criteria. | IQ<70; chronic medical illness; daily prescription medication; head injury with loss of consciousness; traumatic brain injury; neurological disorder; schizophrenia; anorexia nervosa; pervasive developmental disorder; obsessive compulsive disorder; bipolar I disorder or mania; birth weight under 5 lbs.; or severe prenatal (eg, fetal alcohol and/or drug exposure) or perinatal complications (eg, NICU stay); current or lifetime nicotine dependence/ alcohol/substance use disorder; contraindications for safe MRI scan; and Axis I disorder or report of maltreatment that warranted CPS investigation in non-maltreated controls |
| Mannheim | Women aged 18-65 years, PTSD after childhood sexual or physical abuse before the age of 18 years , Sexual or physical assault must be the index trauma, At least 3 criteria of BPD (including criterion 6: affective instability; IPDE), Commitment and possibility to attend weekly therapy sessions for one year; no planned absence for more than 4 weeks in this period. TC: Women aged 18-65 years, Childhood sexual or physical abuse before the age of 18 years. HC: Women aged 18-65 years | PTSD: General exclusion criteria were traumatic brain injuries, current and lifetime schizophrenia or bipolar-I disorder, mental retardation, severe psychopathology or somatic illness that needs to be treated immediately in another setting (e.g., BMI<16), medical conditions making exposure-based treatment impossible, a suicide attempt within the last two months, and substance dependency with no abstinence within two months prior to the study. TC&HC: any current or previous mental disorder, any psychotherapeutic experience or any intake of psychotropic medication lifetime and at the moment. For the current fMRI study, further exclusion criteria were metal implants, pregnancy, left-handedness, and claustrophobia |
| Phan | All: 18-55 years of age, discharge from active military service (post OEF/OIF deployment), ability to read and speak English; PTSD: current PTSD related to combat trauma, CAPS ≥ 40 with greater than 1 month duration of symptoms, CES score ≥ 17; Combat control: CAPS < 20, CES score ≥ 17; | All: life history of bipolar disorder, schizophrenia, mental retardation, passive developmental disorder; severe depressive symptoms as indicated by HAM-D score of ≥ 25; current alcohol/drug dependence within the past 6 months; current suicidal/homicidal ideation; ongoing psychotherapy treatment; left-handedness ; PTSD: presence of clinically significant medical condition or taking a medication which interferes with metabolism of paroxetine; history of hypersensitivity to paroxetine/SSRI; prior failure of response to paroxetine/SSRI for PTSD; history of PTSD or partial/subthreshold PTSD related to a prior deployment or a prior trauma |

**References**

Beck AT, S. R., & Brown GK. (1996). Manual for the Beck Depression Inventory-II. In. San Antonio, TX: Psychological Corporation.

Carleton, R. N., Thibodeau, M. A., Teale, M. J., Welch, P. G., Abrams, M. P., Robinson, T., & Asmundson, G. J. (2013). The center for epidemiologic studies depression scale: a review with a theoretical and empirical examination of item content and factor structure. *PLoS One, 8*(3), e58067. doi:10.1371/journal.pone.0058067

Davidson, J. R., Book, S. W., Colket, J. T., Tupler, L. A., Roth, S., David, D., . . . Feldman, M. E. (1997). Assessment of a new self-rating scale for post-traumatic stress disorder. *Psychol Med, 27*(1), 153-160. doi:10.1017/s0033291796004229

Fortin, J. P., Cullen, N., Sheline, Y. I., Taylor, W. D., Aselcioglu, I., Cook, P. A., . . . Shinohara, R. T. (2018). Harmonization of cortical thickness measurements across scanners and sites. *Neuroimage, 167*, 104-120. doi:10.1016/j.neuroimage.2017.11.024

Hamilton, M. (1960). A rating scale for depression. *J Neurol Neurosurg Psychiatry, 23*, 56-62. doi:10.1136/jnnp.23.1.56

He, Y., Chen, Z. J., & Evans, A. C. (2007). Small-world anatomical networks in the human brain revealed by cortical thickness from MRI. *Cereb Cortex, 17*(10), 2407-2419. doi:10.1093/cercor/bhl149

Kovacs, M. (1985). The Children's Depression, Inventory (CDI). *Psychopharmacol Bull, 21*(4), 995-998.

Lovibond, P. F., & Lovibond, S. H. (1995). The structure of negative emotional states: comparison of the Depression Anxiety Stress Scales (DASS) with the Beck Depression and Anxiety Inventories. *Behav Res Ther, 33*(3), 335-343. doi:10.1016/0005-7967(94)00075-u

Sun, D., Haswell, C. C., Morey, R. A., & De Bellis, M. D. (2018). Brain structural covariance network centrality in maltreated youth with PTSD and in maltreated youth resilient to PTSD. *Dev Psychopathol*, 1-15. doi:10.1017/S0954579418000093

Sun, D., Peverill, M. R., Swanson, C. S., McLaughlin, K. A., & Morey, R. A. (2018). Structural covariance network centrality in maltreated youth with posttraumatic stress disorder. *J Psychiatr Res, 98*, 70-77. doi:10.1016/j.jpsychires.2017.12.015

Teicher, M. H., Anderson, C. M., Ohashi, K., & Polcari, A. (2014). Childhood maltreatment: altered network centrality of cingulate, precuneus, temporal pole and insula. *Biol Psychiatry, 76*(4), 297-305. doi:10.1016/j.biopsych.2013.09.016

Weathers, F. W., Bovin, M. J., Lee, D. J., Sloan, D. M., Schnurr, P. P., Kaloupek, D. G., . . . Marx, B. P. (2018). The Clinician-Administered PTSD Scale for DSM-5 (CAPS-5): Development and initial psychometric evaluation in military veterans. *Psychol Assess, 30*(3), 383-395. doi:10.1037/pas0000486

Weathers, F. W., Keane, T. M., & Davidson, J. R. (2001). Clinician-administered PTSD scale: a review of the first ten years of research. *Depress Anxiety, 13*(3), 132-156. doi:10.1002/da.1029

**Legend for Supplementary Figure 1**

This histogram shows the number of PTSD subjects and controls in each age group. Age groups with highest number of PTSD subjects is 30-40 years (n=375), followed by 20-30 years (n=347). Age groups with highest number of control subjects is 20-30 years (n=550), followed by 30-40 years (n=420).
