## Supplementary figures and images for "Structural Covariance Networks in Post-Traumatic Stress Disorder: A Multisite ENIGMA-PGC Study"

### Supplemental Figure 1

#### Supplemental Figure 1

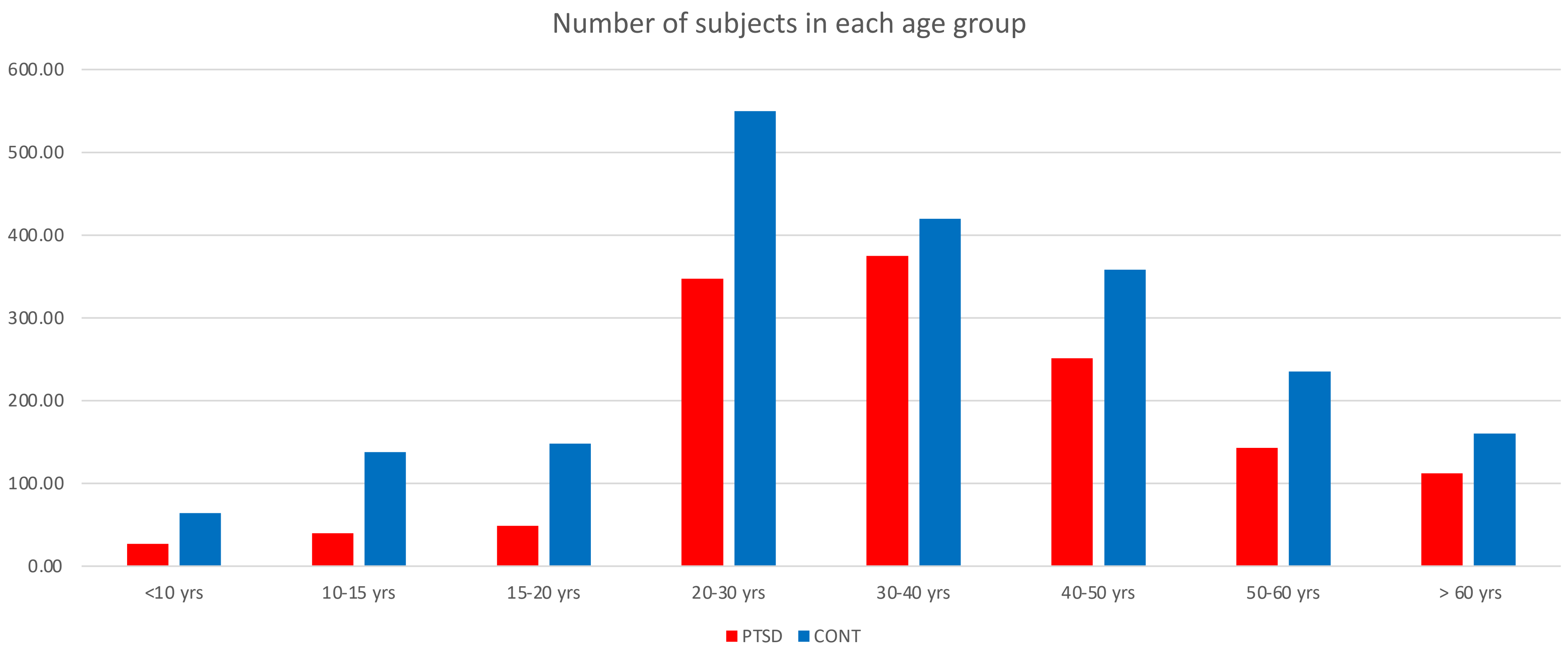
